# Adaptation to ER Stress by Slt2, Counterpart of Human MAP Kinase ERK1/2, via Enhancing Splicing and Translation of *HAC1* mRNA in *Saccharomyces cerevisiae*

**DOI:** 10.1101/2023.11.19.567283

**Authors:** Jagadeesh Kumar Uppala, Anish Chakraborty, Jasmine George, Kimberly Ann Mayer, Chandrima Ghosh, Ritisha Dey, Pradeep Chaluvally-Raghavan, Madhusudan Dey

**Affiliations:** Department of Biological Sciences, UW-Milwaukee, Milwaukee, WI-53211; Department of Obstetrics and Gynecology, Medical College of Wisconsin, Milwaukee, WI-53226

**Author notes:** Corresponding author: Madhusudan Dey, Department of Biological Sciences, University of Wisconsin-Milwaukee 3209 N Maryland Ave, Milwaukee, WI-53211.

**Keywords:** Protein kinase, Endoplasmic reticulum stress (ER stress), Unfolded protein response (UPR), Mitogen-activated protein kinase (MAPK), Extracellular-signal-regulated kinase (ERK), Ire1, Slt2, and HAC1 mRNA

## Abstract

Unfolded protein response (UPR) is a cellular strategy to increase the protein folding capacity of cells in response to stress within the endoplasmic reticulum (ER). In metazoan cells, three major UPR sensors Ire1, PERK and ATF6 work in concert by simultaneously activating intracellular signaling pathways and modulating a series of physiological processes such as attenuation of the general protein synthesis and expression of protein chaperones. In yeast *Saccharomyces cerevisiae*, Ire1 is known to be the only UPR sensor, which mediates splicing of *HAC1* mRNA in the cytoplasm and derepresses its translation. Hac1 is a transcription factor that increases the expression of protein folding enzymes and chaperones, thus enhancing the protein folding capacity of cells. In this study, we provide compelling evidence that kinase Slt2 plays a significant role in facilitating both the splicing and translation of *HAC1* mRNA, while also serving as a key mediator in the activation of UPR genes through an alternative route. We also provide evidence that human extracellular signal-regulated kinase 1 (ERK1) or ERK2 served as a functional substitute for yeast Slt2 in the context of UPR. Furthermore, ERK1 exhibits an enhanced activation in human primary cells when grown in the presence of ER stressor. These findings collectively suggest that Slt2 responds to ER stress by activating the Ire1 pathway as well as initiating a parallel signaling pathway.

## Introduction

Mitogen (growth factor)-activated protein kinases (MAPKs) are group of kinases, which are activated in response to a diverse set of extracellular and intracellular stimuli including growth factors, cytokines, and a variety of extra- and intra-cellular stresses (1,2). Following stimulation, three MAP kinases are typically activated sequentially: a proximal MAP kinase kinase kinase (MAPKKK or MAP3K) which phosphorylates and activates a MAP kinase kinase (MAPKK or MAP2K), which then phosphorylates and activates a terminal MAP kinase (MAPK). These three MAPKs constitute a core 3-tier MAPK module with a scaffold protein, which is known to embed within the signaling circuits and regulate a diverse set of cellular functions, ranging from cell growth, proliferation, differentiation and programmed cell death(3).

In humans, there are at least four major groups of MAPKs: (1) extracellular signal-regulated kinase (ERK) with two identical isoforms ERK1 and ERK2(4,5), big MAP kinase 1 (BMK1), also called ERK5(6), containing a transcription activation domain (TAD), (3) Jun N-terminal kinase (JNK) with three isoforms JNK1, JNK2, and JNK3(7–9), and (4) the p38 kinase with four isoforms p38α, p38β, p38γ and p38δ(10–12). Each MAPK module is stimulated by a unique set of signal transduction pathways. For instance, the ERK family members are stimulated primarily by exposure of cells to phorbol esters, growth factors and/or cytokines. Following stimulation, ERK1/2 are activated upon phosphorylation by the upstream kinases MEK1 and MEK2, which are simultaneously activated upon phosphorylation by Raf kinases (A-Raf, B-Raf and Raf1). Like ERK1, ERK5 is activated upon phosphorylation by upstream kinases MEK5α and MEK5β that are activated upon phosphorylation by kinases MEKK2 or MEKK3(13). In contrast, the p38 and JNK family members respond primarily to exposure of cells to intracellular and extracellular stressful stimuli, thus known as the stress activated protein kinases (SAPK) (14). After stimulation, JNK and its isoforms are activated by upstream kinases MKK4 and MKK7(15), whereas p38 and its isoforms are activated by kinases MKK3 and MKK6(16).

In the budding yeast *Saccharomyces cerevisiae*, five distinct MAPKs (Fus3, Kss1, Hog1, Slt2 and Smk1) are known to mediate the cellular signals for various cellular processes(17) (**Fig 1A**). Among these MAPKs, the Hog1 and Slt2 kinases are known to be stress activated protein kinases (SAPKs). The Hog1 kinase is the yeast orthologue of mammalian kinase p38(11,18), which is controlled by two MAP3K kinases (Ste11 and Ssk2) and one MAP2K (Pbs2) (**Fig 1A**). Additionally, the catalytic activity of the Hog kinases is directed by a scaffold protein Ste5. This HOG pathway is known to be primarily activated when the osmolarity of the cellular environment changes rapidly(18). The MAP kinase Slt2 is generally regarded as the yeast orthologue of human ERK5 (19) and activated under a variety of stressful conditions such as high temperature(20) hypo-osmotic shock (21) and polarized growth(22). The kinase function of Slt2 is controlled by MAP3K Bck1 and two redundant MAP2Ks (Mkk1 and Mkk2) (22,23). The active Slt2 MAPK module integrates with specific signaling molecules to reduce the cell wall stress (17,24). Recently, it has also been shown that the Slt2 MAPK module becomes active when the protein folding functions of the endoplasmic reticulum (ER) go awry(25). However, the molecular function of the Slt2 kinase in ER function is still not clear.

**Figure 1:**
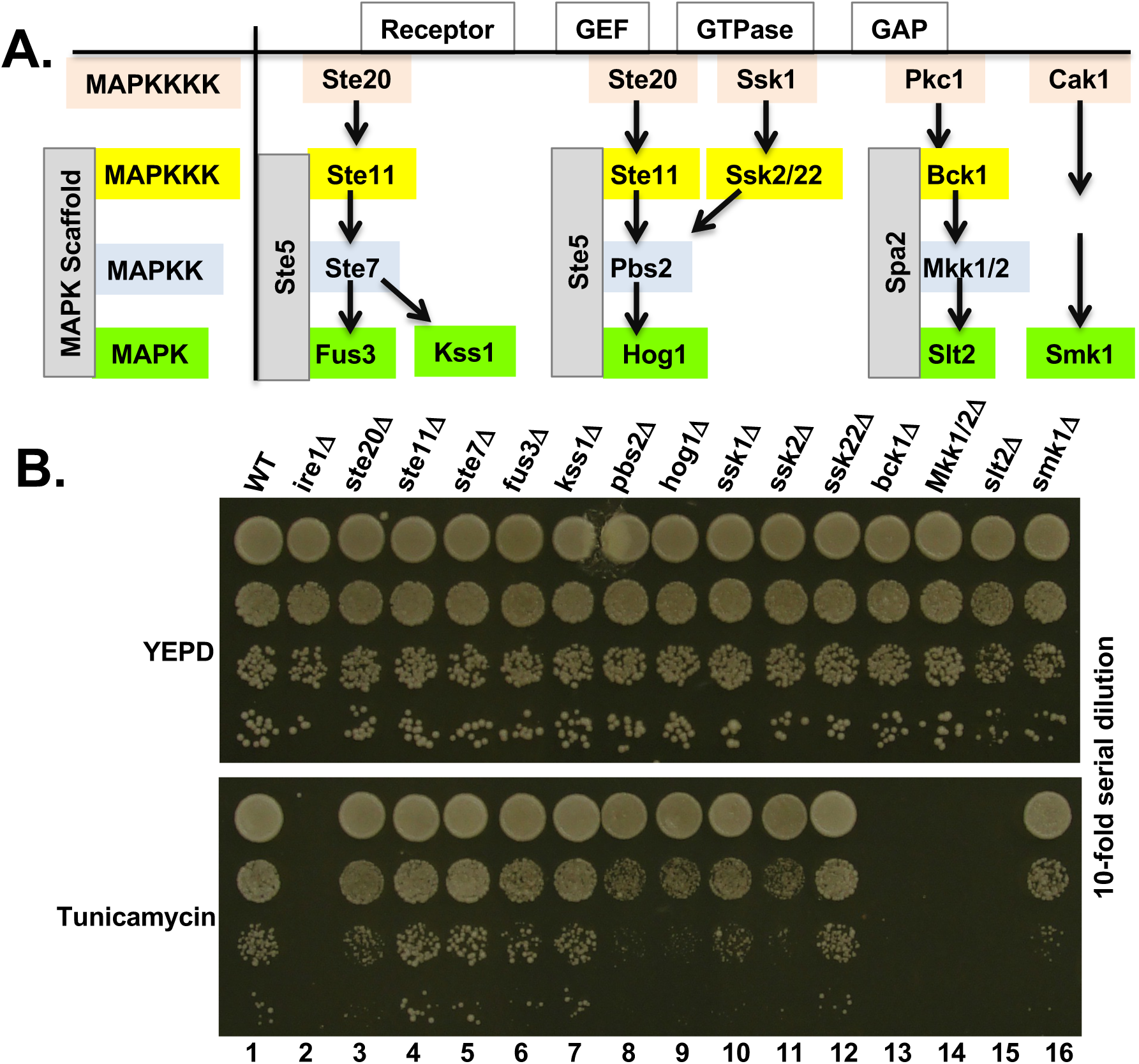
Yeast cells lacking the MAP kinase Slt2 is sensitive to tunicamycin. **(A)** A simplified cartoon representation of the major MAP kinase signaling pathways in yeast *Saccharomyces cerevisiae*. **(B)** The indicated yeast deletion strains were serially diluted, spotted and grown on the YEPD and the same medium containing tunicamycin at 30°C for 48 hours.

The ER is an intracellular organelle where most secretory and membrane proteins fold and mature before traveling onward to their destination. In cases when the loading of newly synthesized proteins exceeds the folding capacity of ER, unfolded proteins accumulate inside the ER lumen, causing an imbalance in the ER protein homeostasis, a condition known as “ER stress”. ER stress then triggers the unfolded protein response, an adaptive strategy to restore the ER protein homeostasis controlled by three major ER-resident proteins Ire1 (a dual kinase RNase) (26–28), PERK (a Ser/Thr protein kinase)(29) and ATF6 (a transcription factor)(30). These proteins work in concert by simultaneously activating parallel intracellular signaling pathways and modulating a series of physiological processes such as attenuation of the general protein synthesis, expression of the ER-resident protein chaperones(31–33) and the activation of the ER-assisted degradation (ERAD) of unfolded proteins(34–36).

Ire1 is a conserved sensor from yeast to humans. During ER stress, Ire1 initiates the UPR by cleaving the *HAC1* mRNA in yeast cells or *XBP1* mRNA in metazoan cells. The cleaved *HAC1* mRNAs are ligated by tRNA ligase(37), whereas *XBP1* mRNAs by RtcB(38). The spliced *HAC1* or *XBP1* mRNA yields a transcription factor that activates several genes involved in accelerating the protein folding capacity of ER. Along with the Ire1 pathway, metazoan cells simultaneously activate the PERK and ATF6 pathways. The active PERK phosphorylates the translation initiation factor eIF2α. The phosphorylated eIF2α activates the expression of the transcription factor ATF4 utilizing two short open reading frames (ORFs) present in its leader sequence(39). ATF4 activates expression of several ER-resident enzymes and protein modifiers. Unlike Ire1 and PERK, ATF6 itself is a transcription factor and processed uniquely for its activation. ATF6 trans-locates from the ER to Golgi complex where it is site-specifically cleaved to release its active transcription activation domain. Collectively, Hac1/Xbp1, ATF4 and ATF6 activate the expression of ER-resident enzyme genes and chaperones, which accelerate the protein folding capacity of cells and clear the unfolded proteins, which eventually restore the ER protein homeostasis (adaptive phase of UPR). However, under conditions of severe or prolonged ER stress, the adaptive program of UPR integrates with the apoptotic program and executes cell death (terminal phase of UPR). For example, Ire1 broadens its substrate specificity and non-specifically cleaves mRNAs associated with the ER in a process known as the regulated Ire1 dependent degradation of RNA or RIDD that could promote cell death unfolded proteins (40,41). Like Ire1, PERK activates the transcriptional regulator CHOP that in turn activates apoptosis by increasing the expression of BIM and DR5(42) Similarly, ATF6 mediates apoptosis either by activating CHOP and/or by specific suppression of Mcl-1(43).

The yeast MAP kinase Slt2 is known to phosphorylate and activate several proteins, including the transcription factor Rlm1 and Swi4, which are important for expression of genes involved in cell wall biosynthesis(44,45). Slt2 has also been reported to play important role in both adaptive and apoptotic responses of UPR (46–48); however, less is known about Slt2’s molecular role in ER stress response. In this study, we show that Slt2 promotes Ire1-mediated ER stress response and plays a crucial role in mitigating the prolonged ER stress that might not be resolved by immediate activation of Ire1-mediated UPR. We further show that the human orthologs ERK1 and ERK2 can complement the Slt2 function in alleviating the ER stress.

## Results

### 1. MAP kinase Slt2 promotes Ire1-mediated ER stress response

To elucidate the involvement of MAP kinases in the yeast unfolded protein response, we compared growths between wild-type (WT) and its isogenic strains lacking a single MAP3K, MAP2K or MAPK on the medium containing tunicamycin, a chemical inducer of ER stress. Yeast cells lacking the kinase Ste20 (MAP4K), Ste11 (MAP3K), Ste7 (MAP2K), Fus3 (MAPK) or Kss1 (MAPK) were resistant to tunicamycin at concentration 0.25 μg/ml (**Fig 1B**, rows 3-7). At the same concentration of tunicamycin, yeast cells lacking the kinase Ssk1 (MAP4K), Ssk2 (MAP3K), Pbs2(MAP2K) or Hog1 (MAPK) were mildly sensitive to tunicamycin (**Fig 1B**, rows 8-11, whereas yeast cells lacking the kinase Bck1 (MAP3K), Mkk1/2 (MAP2K) and Slt2 (MAPK) were severely sensitive to tunicamycin (**Fig 1B**, rows 13-15), like an *ire1Δ* cell (**Fig 1B**, row 2). Tunicamycin prevents protein maturation by inhibiting the co-translational N-linked glycosylation of the nascent polypeptides, resulting in accumulation of unfolded proteins inside the ER lumen. Thus, the sensitivity to tunicamycin indicates that, in addition to the classic UPR pathway, at least two other pathways to be involved in the response to ER stress, including the Slt2 MAPK pathway and the osmo-sensing HOG1 MAPK pathway.

The above findings align perfectly with the earlier studies conducted by Chen et al. (2005), which showed that cells lacking the components of MAP kinase Slt2 pathway display severe sensitivity to tunicamycin (46). Chen et al. (2005) also showed that the levels of Hac1 protein expression were comparable in wild-type (WT) and *slt2Δ* strains when cell were exposed to the ER stressor tunicamycin for an hour in a liquid culture. These results led them to conclude that the Slt2 pathway independently controls the ER stress response, which is distinct from the Ire1 pathway. However, we posit that a one-hour tunicamycin treatment might not be adequate to fully elicit the ER stress response and replicate the sensitivity to tunicamycin observed on the solid medium (**Fig 1B**, rows 13-15). Consequently, we prolonged the tunicamycin treatment for a minimum of 2 hours and investigated the involvement of MAP kinases in the Ire1-mediated ER stress response by comparing the expressions of Hac1 protein in the MAP kinase deletion strains that exhibited mild or severe sensitivity to tunicamycin.

The yeast strains were grown in a liquid YEPD medium until the OD_600_ value reached to 0.5-0.6. Subsequently, cells were treated with 1 μg/ml tunicamycin for 2 hours. Tunicamycin-treated cells were harvested and whole cell extracts were prepared. Whole cell extracts were utilized for Western blot analysis to examine the expression of Hac1 protein, while Reverse Transcriptase (RT)-PCR analysis was conducted to assess the splicing of *HAC1* mRNA (see **Materials and Methods**). As shown in **Fig 2A**, the amount of Hac1 protein was reduced in cells lacking Bck1 (by 23.5%), Slt2 (by 68.2%) and Mkk1/2 (by 47.8%). The *HAC1* mRNA splicing was reduced in cells lacking Bck1 (by 22.8%), Slt2 (by 46.7 %) and Mkk1/2 (by 14.6%) compared to their isogenic wild type cells (**Fig 2B**). Furthermore, we monitored the Hac1 protein expression in both WT and *slt2Δ* strains grown in the presence and absence of an ER stressor DTT (5 mM), a reducing agent recognized for disrupting disulfide bonds. Consistently, we observed that Hac1 protein expression was reduced (68%) in the *slt2Δ* strain compared to its isogenic WT strain (**Fig 2C**). The significant reductions in splicing of *HAC1* mRNA and expression of Hac1 protein, together with severe sensitivity to tunicamycin at a low concentration (0.2 μg/ml), suggest that the MAP kinase Slt2 contributes to folding of ER client proteins by directly influencing the Ire1-mediated *HAC1* mRNA splicing and translation from the spliced mRNA.

**Figure 2:**
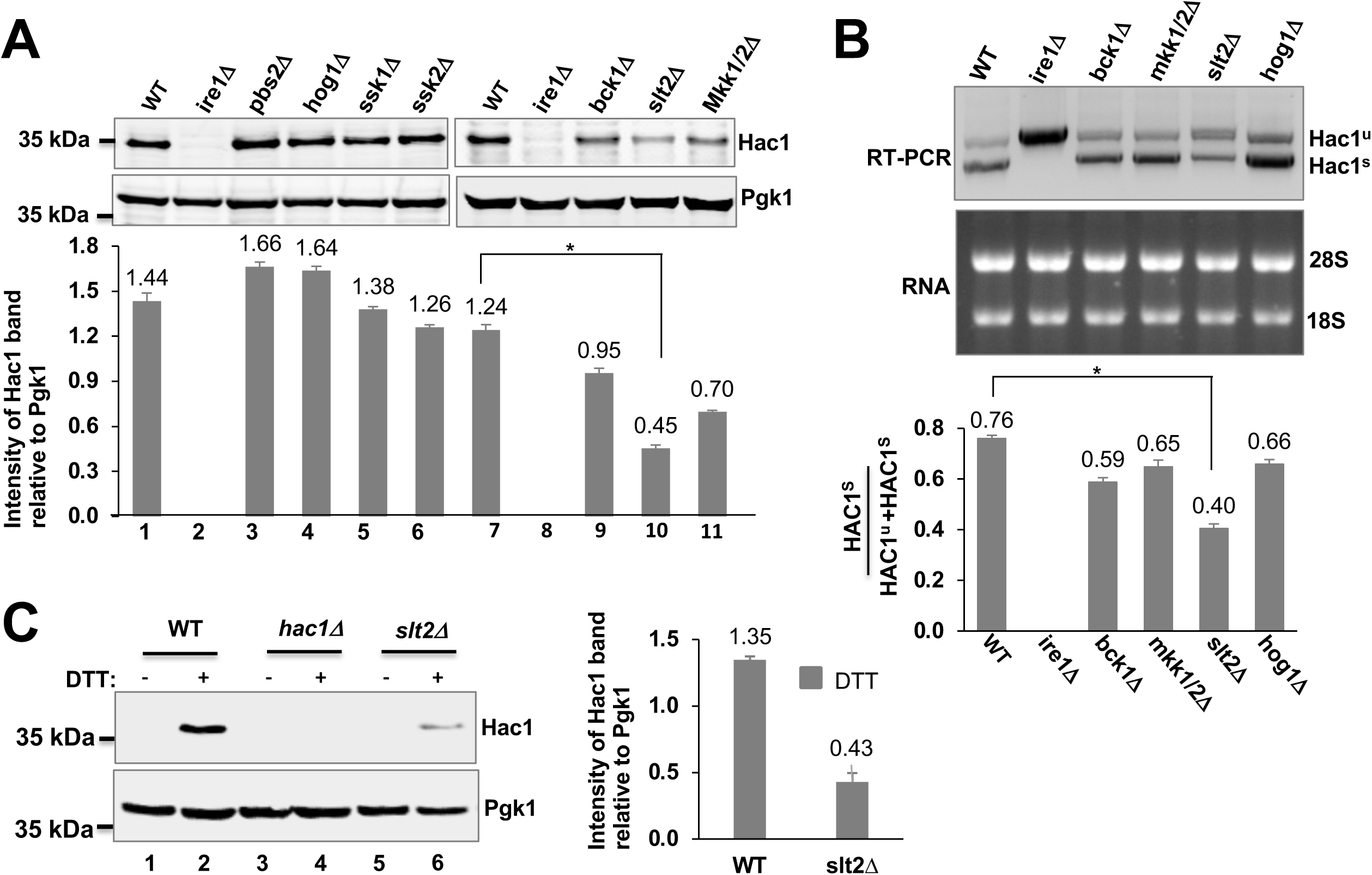
Reduced splicing and translation of *HAC1* mRNA in the *slt2Δ* strain. **(A)** (Upper panel) The indicated yeast deletion strains were grown in the YEPD medium until they reached the OD_600_ value between 0.5 and 0.6. Cells were then exposed to tunicamycin (1μg/ml) and harvested after 2 hours. Whole cell extracts (WCEs) were prepared and subjected to Western blot analysis using antibodies specific to Hac1 and Pgk1 proteins. (Lower panel) The intensities of Hac1 and Pgk1 protein bands were quantified using the ImageJ software. The bar diagram depicts the relative intensities of Hac1 protein bands. **(B)** The indicated yeast deletion strains were grown in the YEPD medium until they reached the OD_600_ value ∼0.5-0.6. Cells were then exposed to tunicamycin (1μg/ml) and harvested after 2 hours. Total RNA was extracted and subjected to RT-PCR analysis to distinguish between the spliced (HAC1^s^) and un-spliced (HAC1^u^) *HAC1* mRNAs. (Lower panel) The intensities of HAC1^s^ mRNAs were quantified using the ImageJ software. The bar diagram depicts the relative intensities of HAC1^s^ mRNAs. **(C)** The indicated yeast deletion strains were grown in the YEPD medium in the absence and presence of DTT. WCEs were prepared and subjected to Western blot analysis using antibodies specific to Hac1 and Pgk1 proteins. The intensities of Hac1 and Pgk1 protein bands were quantified using the ImageJ software. (Right panel) The bar diagram depicts the relative intensities of Hac1 protein bands.

To further test our results that Slt2 kinase contributes to translation from the spliced *HAC1* mRNA, we investigated the Hac1 protein expression from an intron-less *HAC1* mRNA in the *ire1Δ* strain. The intronic sequence of the *HAC1* ORF was removed to make a plasmid that could express an intron-less *HAC1* mRNA constitutively. This engineered plasmid was designated as the HAC1-constitutive (HAC1^c^) (**Fig 3A**). The HAC1^c^ plasmid was introduced in the strain lacking the kinase Ire1 or Slt2. The resultant strains were tested for their sensitivity to tunicamycin (**Fig 3B**) as well as their ability to produce Hac1 protein (**Fig 3C**). As anticipated, the *ire1Δ* strain containing the HAC1^c^ plasmid grew on the medium containing tunicamycin (**Fig 3B**, row 2). The tunicamycin-resistant phenotype was corelated with the Hac1 protein expression (**Fig 3C**, lane 1). In contrast, the *slt2Δ* strain harboring the HAC1^c^ plasmid did not grow on the tunicamycin medium (**Fig 3B**, row 4). Interestingly, we observed the presence of Hac1 protein in the *slt2Δ* strain (**Fig 3C**, lane 4), albeit an expression level 3-fold lower when compared to the *ire1Δ* strain (**Fig 3C**, lower panel). These data suggest that Slt2 kinase contributes to translation from the spliced *HAC1* mRNA and Hac1 protein alone was not sufficient to promote cell growth under conditions of ER stress.

**Figure 3:**
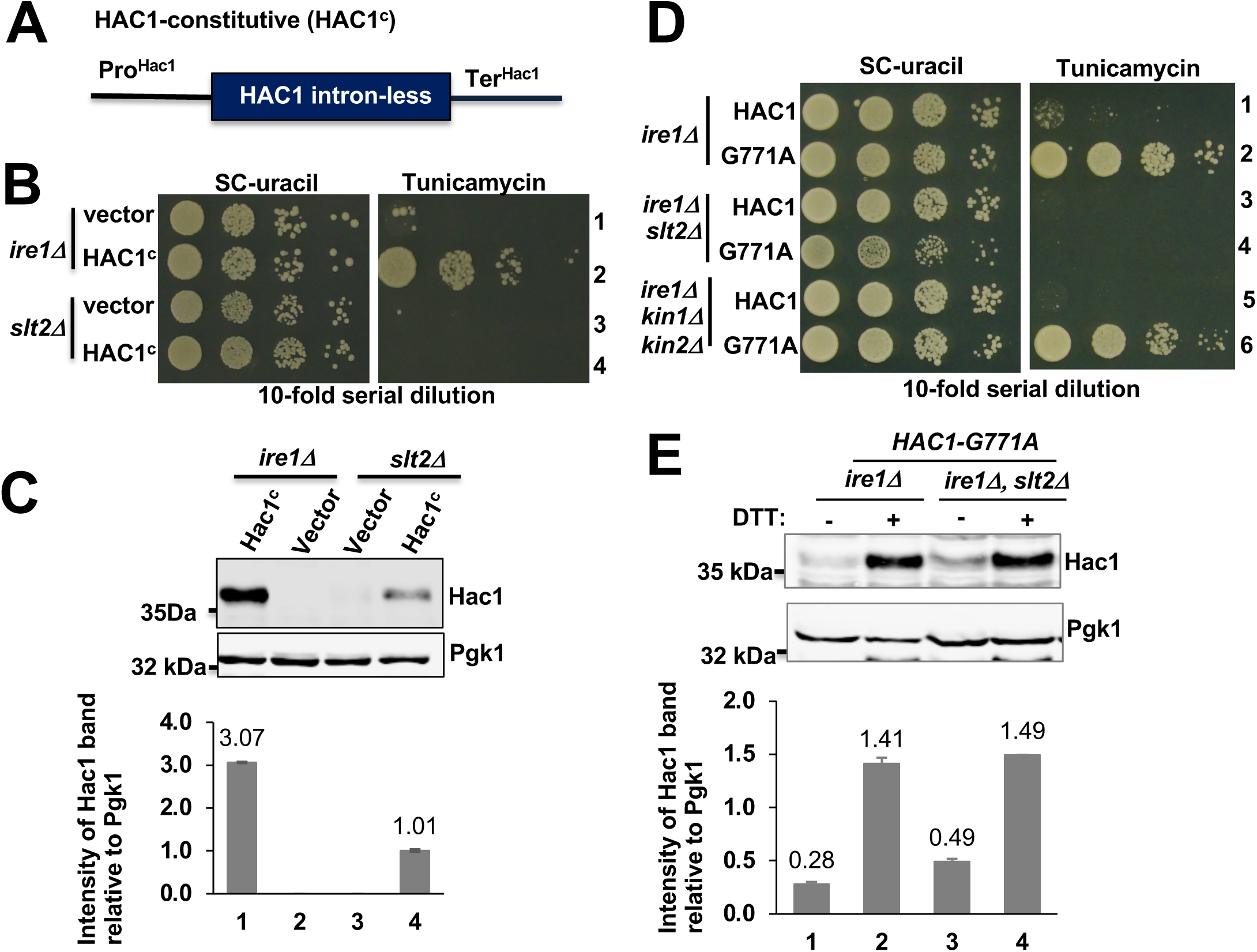
Reduced translation of an intron-less *HAC1* mRNA in the *slt2Δ* strain. **(A)** Schematic diagram of the constitutive HAC1 allele (HAC1^c^). The intronic sequence from the *HAC1* open reading frame was removed. **(B)** The indicated yeast strains containing a vector plasmid or the same vector bearing a constitutive HAC1 allele (HAC1^c^) were grown, serially diluted, spotted and grown on the YEPD and the same medium containing tunicamycin at 30°C for 48 hours. **(C)** WCEs were prepared from the indicated yeast strains as shown in the panel (**B**). WCEs were subjected to Western blot analysis using antibodies specific to Hac1 and Pgk1 proteins. The intensities of Hac1 and Pgk1 protein bands were quantified using the ImageJ software. (Lower panel) The bar diagram depicts the relative intensities of Hac1 protein bands. **(D)** The indicated yeast strains expressing the wild-type HAC1 or HAC1-G771A allele were grown, serially diluted, spotted and grown on the YEPD and the same medium containing tunicamycin at 30°C for 48 hours. **(E)** WCEs were prepared from the indicated yeast strains expressing the HAC1-G771A allele. WCEs were subjected to Western blot analysis using antibodies specific to Hac1 and Pgk1 proteins. The intensities of Hac1 and Pgk1 protein bands were quantified using the ImageJ software. (Lower panel) The bar diagram depicts the relative intensities of Hac1 protein bands.

To further confirm that the tunicamycin-resistant phenotype associated with the ER stress response was not solely due to expression of Hac1 protein, we examined the expression of Hac1 protein from the translationally de-repressed *HAC1-G771A* mRNA(49). We also assessed its influence on tunicamycin sensitivity. As anticipated, the *ire1Δ* and *ire1Δ kin1Δ kin2Δ* strains containing the *HAC1-G771A* allele grew on the medium containing tunicamycin (**Fig 3D**, rows 2 and 6) and this growth was corelated the Hac1 protein expression from the un-spliced mRNA (**Fig 3E**, lane 2). Consistent with the previous result(49), we also observed that Hac1 expression from the un-spliced *HAC1-G771A* mRNA was increased when cells were treated with ER stressor DTT (**Fig 3E**). Interesting, we observed that the *ire1Δslt2Δ* strain harboring the *HAC1-G771A* allele was unable to grow on the tunicamycin medium (**Fig 3D**, row 4), even though a significant amount of Hac1 protein expression was detected when cells were grown in the absence or presence of DTT (**Fig 3E**, lanes 3 and 4). These data confirmed that Hac1 protein alone was insufficient to initiate the ER stress response.

### 2. MAP kinase Slt2 is maximally activated during the adaptive stage of UPR

To further investigate the role of Slt2 in the Hac1-mediated ER stress response, we expressed extra-copies of Slt2 protein from a low copy-numbered centromeric (CEN4) plasmid in the wild-type cells and monitored the Hac1 expression. When Slt2 was expressed from a CEN4 plasmid, it complemented the loss of Slt2 and restored growth on the tunicamycin medium (**Fig 4A**, compare rows 3 and 4). When the wild type cell contains a *CEN4* plasmid bearing the *SLT2* gene, the expression of Slt2 protein was enhanced ∼3-fold (**Fig 4B**, compare lanes 1 and 2). A gradual increase of the Hac1 protein expression was observed in the wild-type cell with a vector plasmid (**Fig 4C**, compare lanes 3, 4 and 5). Interestingly, we observed that the expression of Hac1 protein in the wild-type cell containing the *SLT2* gene was enhanced following 1 hour of DTT treatment (**Fig 4C**, compare lane 1 and 4). These results suggest that MAP kinase Slt2 plays a significant role in the cellular response to ER stress mediated by Hac1. Furthermore, we compared growths of *ire1Δ*, *slt2Δ* and *ire1Δslt2Δ* strains on the medium containing tunicamycin at concentration 0.2 μg/ml. Consistent with the previous report (ref), we found that the *ire1Δslt2Δ* double deletion strain was more sensitive to tunicamycin compared to the single deletion strain (data not shown). These results collectively suggest that the MAP kinase Slt2 plays a significant role in the cellular response to ER stress not only by activating the Ire1 pathway but also by initiating a parallel signaling pathways.

**Figure 4:**
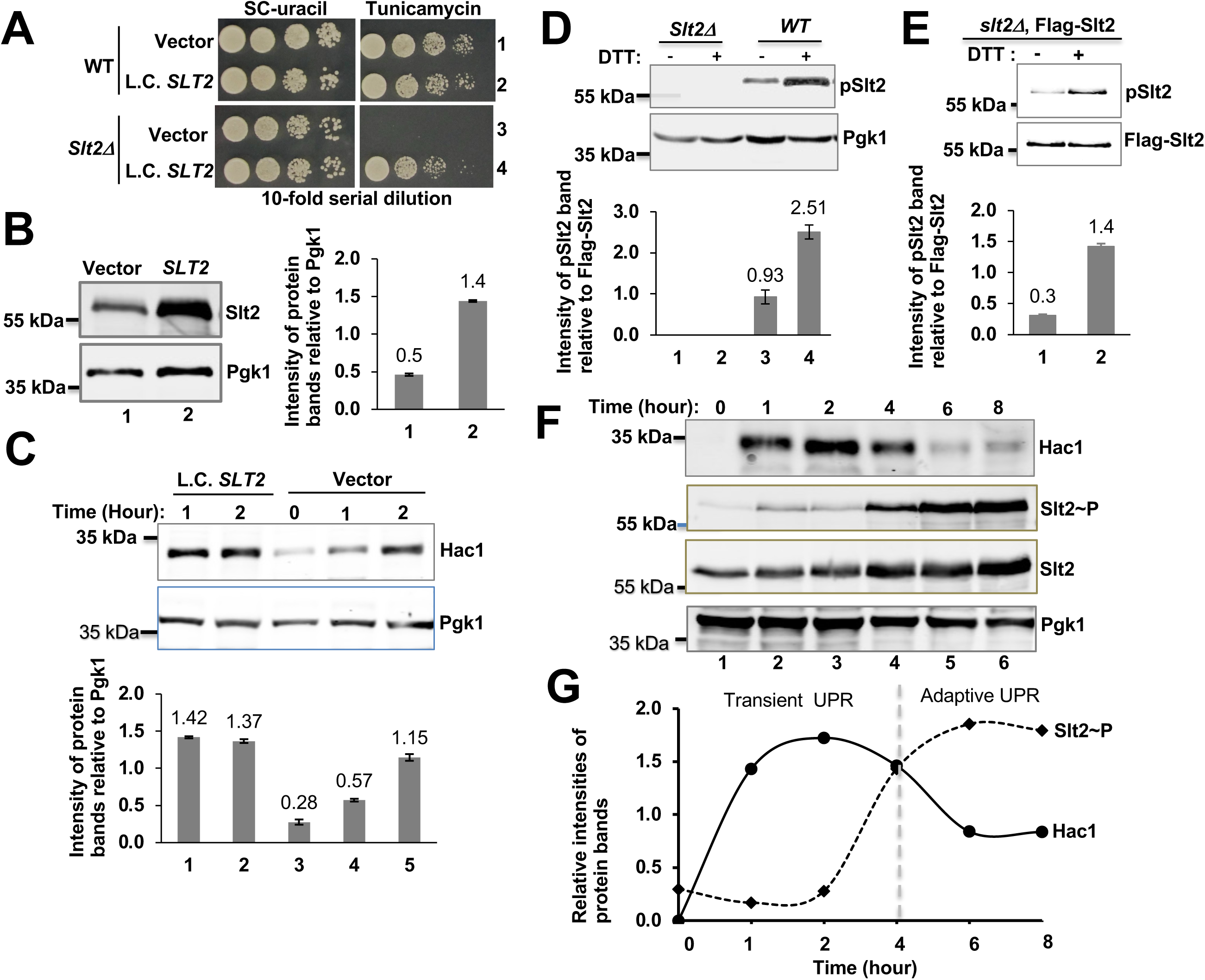
Activation of Slt2 kinase during the adaptive stage of ER stress. **(A)** The *slt2Δ* strains bearing a low copy numbered (L.C.) plasmid or the same plasmid bearing the *SLT2* gene were tested for growth on the SC-uracil medium or the same medium containing tunicamycin. **(B)** WCEs were prepared from the yeast strains shown in the **Fig 4A** (rows 1 and 2) and subjected to Western blot analysis using antibodies specific to Slt2 and Pgk1. (Right, lower panel) The bar diagram depicts the relative intensities of Slt2 and Pgk1 protein bands. **(C)** The yeast strains bearing a low copy numbered (L.C.) plasmid or the same plasmid bearing the *SLT2* gene were grown in the presence of DTT for the indicated time. WCEs were prepared and subjected to Western blot analysis using antibodies specific to Hac1 and Pgk1 proteins. (Lower panel) The bar diagram depicts the relative intensities of Hac1 and Pgk1 protein bands obtained from the Fig 4C. **(D)** The indicated yeast strains were grown in the absence (-) and presence (+) of DTT. WCEs were prepared and subjected to Western blot analysis using antibodies specific to phosphorylated Slt2 (pSlt2) or Pgk1. **(E)** The *slt2Δ* strain expressing a Flag-epitope tagged Slt2 (Flag-Slt2) under the control of a GAL1 promoter was grown in the absence (-) and presence (+) of DTT. WCEs were prepared and subjected to Western blot analysis using antibodies specific to phosphorylated Slt2 (pSlt2) or Flag epitope. **(F)** The wild-type yeast strain was grown till the OD600 value reached ∼0.5 to 0.6. Cells were then exposed to DTT. Cells were harvested after 1, 2, 4, 6 and 8 hours. WCEs were prepared and subjected to Western blot analysis using antibodies specific to HAC1, pSlt2, Slt2 and Pgk1. **(G)** The intensities of Hac1, Slt2 and Pgk1 protein bands were quantified using the ImageJ software. The line diagram depicts the relative intensities of Hac1 and Slt2 protein bands relative to Pgk1.

In the active state, typical MAPKs are phosphorylated at the conserved threonine (Thr or T) and tyrosine (Tyr or Y) residues of the TXY motif within their activation loop (X represents any other amino acid) stresses (1,2). Therefore, we monitored the phosphorylation within the activation loop of the Slt2 kinase by Western blot analysis using an antibody specifically raised to detect the phosphorylated activation loop of ERK1 (**Materials and Methods**), thereby detecting the phosphorylated activation loop of Slt2. We initially observed that phosphorylation of Slt2 within its activation loop was enhanced ∼2.5-fold when wild type cells were grown in the presence of DTT (5 mM) for 2 hours compared to cells grown in the absence of DTT (5 mM) for 1 hour (**Fig 4D**, compare lanes 3 and 4). Additionally, we observed that a Flag-tagged Slt2 protein (Flag-Slt2) that complemented the *slt2Δ* strain like WT Slt2 (data not shown) was also phosphorylated almost 4-fold within its activation loop when cells were grown in the presence of DTT (5 mM) for 2 hours (**Fig 4E**, lane 2). These results are consistent with earlier studies that the MAP kinase Slt2 is activated through the phosphorylation of residues T190 and Y192 within its activation loop in response to cell wall stress(ref) and the ER stress (46).

Next, we investigated the phosphorylation pattern within the activation loop of the Slt2 kinase during the induction of ER stress. WT cells were grown until the OD_600_ value reached to ∼0.6, followed by the treatment with DTT (5 mM), and collection of cells after 1, 2, 4, 6 and 8 hours. Whole cell extracts (WCEs) were then prepared and subjected to Western blot analysis using polyclonal antibodies specific for Hac1, Slt2 and Pgk1, alongside a phospho-specific antibody targeting the activation loop of ERK1 (which detects phosphorylated activation loop of Slt2). As expected, the expression of Hac1 protein was observed within 1 hour of DTT treatment (**Fig 4F**, upper panel, lane 2). Interestingly, we observed that Hac1 expression was reduced at 6^th^ and 8^th^ hours after DTT treatment. Remarkably, a marginal increase in the phosphorylation within the activation loop of Slt2 was observed following 1 and 2 hours of DTT-induced stress (**Fig 4F**, middle panel, lane 3). The substantial increase in the Slt2 phosphorylation (19-fold) was observed at and after the 4^th^ hour of DTT treatment (**Fig 4F**, middle panel, lanes 4, 5 and 6). These findings indicate that the cellular response to ER stress involves two distinct stages: an initial transient phase followed by an adaptive phase (**Fig 4G**).

The initial phase (see, transient UPR as shown in **Fig 4G**), which is rapid to mount a survival response, is primarily mediated by the active Ire1, leading to translation of Hac1 protein. Hac1, in turn, activates the expression of protein folding enzymes and chaperones that increase the protein folding capacity of cells. The slower second phase (see, adaptive UPR as shown in **Fig 4G**) involves the activation of both Slt2 and Ire1 proteins. In this slower response, Ire1 pathway is suppressed, leading to reduced expression of Hac1 protein and the restoration of the normal physiological response or adaptation to the ER stress with the help of kinase Slt2. However, if the severe stressful event persists for longer period, cells may enter the apoptotic pathway and execute cell death. These observations help explain the severe sensitivity of *slt2Δ* strain to tunicamycin when grown on a solid medium for extended periods (**Fig 1B**, row 15). Together, our results suggest that Slt2 kinase achieves its maximum activation during the adaptive stage of UPR.

### 3. The predicted helix-**α**L of Slt2 plays an important role in ER stress response

The NCBI BLAST search against human genome database showed that ERK5, ERK1, ERK2 and p38 MAP kinases were close orthologs of Slt2 kinase with sequence identities 48.32% (E value = 8X10^-113^), 49.43% (E value = 3X10^-109^), 49.57% (E value = 2X10^-108^) and 45.83% (E value = 2X10^-99^), respectively (**Fig 5A**). Multiple protein sequence alignment also showed that the kinase domains (KDs) of Slt2 (residues 23-318), ERK1 (residues 42-330), ERK2 (residues 25-313) and ERK5 (residues 54-346) exhibited the highest degree of homology (**Fig 5B**). Additionally, we found that ERK1, ERK2, ERK5 and Slt2 share a common sequence element with 50% homology located at the C-terminal end of their kinase domains. This element adopts a helical structure, known as “helix-αL”, as observed in the crystal structures of ERK1 and ERK2(50). The helix-αL interacts with the helix-αC of the ERK1/ERK2 kinase domain, indicating that it might regulate the kinase domain function.

**Figure 5:**
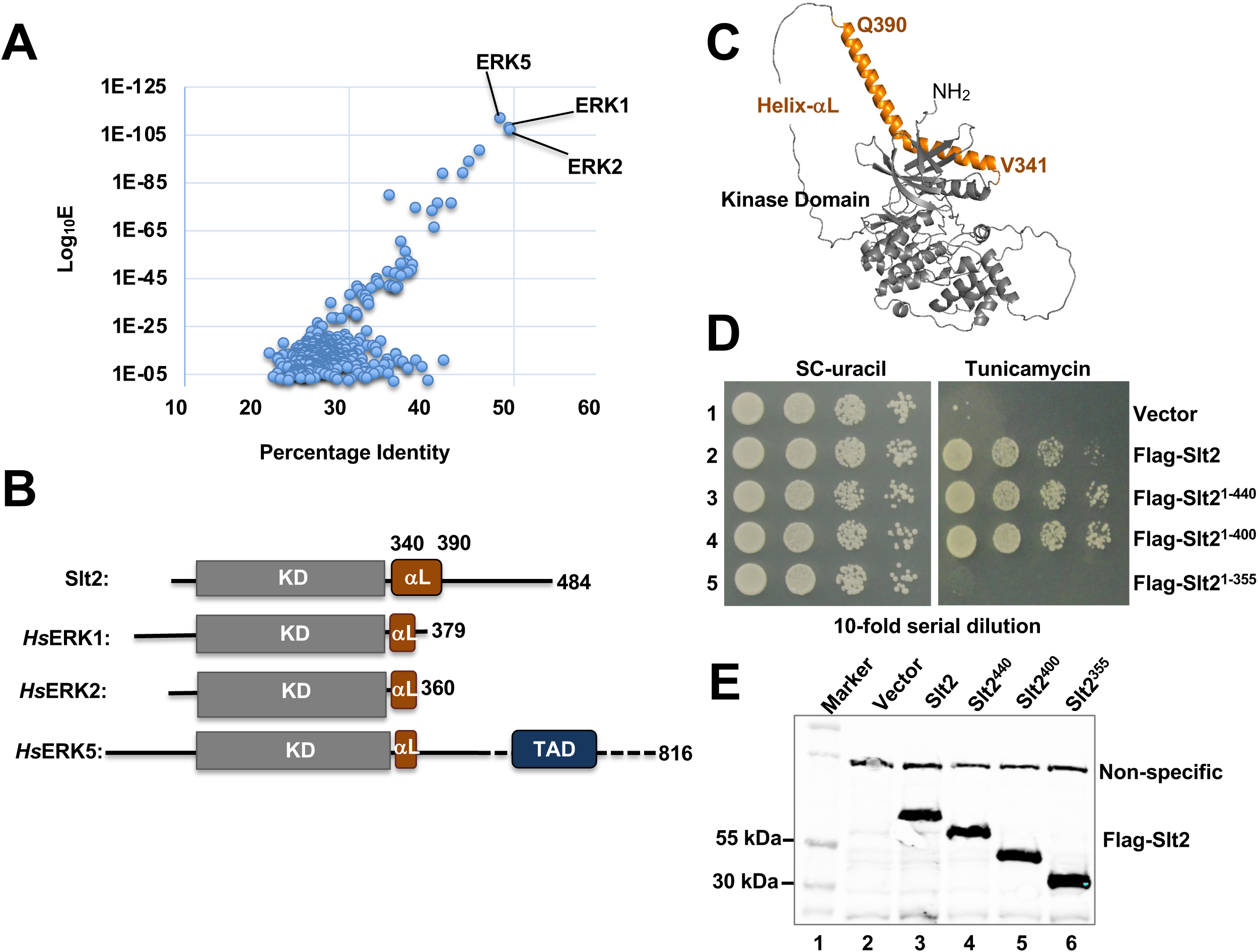
The predicted helix-αL requires for Slt2 function during the ER stress response. **(A)** The Slt2 protein sequence was used as a query in a BLAST search against the NCBI human genome database. The scatter plot displays the percentage identity and their respective E-values. **(B)** The schematic diagram of the indicated MAP kinases and their total number of amino acids content. The kinase domain (KD, grey rectangular boxes), helix-αL (orange boxes) and transcription activation domain (TAD, blue box) are indicated. **(C)** The PyMol software was used to analyze the structure of Slt2 as determined by AlphaFold. The kinase domain is colored in grey while the helix-αL in orange. **(D)** The *slt2Δ* strain containing a Yes2 vector plasmid and the same vector expressing the indicated Flag-tagged Slt2 proteins were tested for growth on the complete synthetic (SC) medium without uracil and the same medium containing tunicamycin. **(E)** WCEs were prepared from yeast strain indicated in the panel (**D**) and subjected to Western blot analysis using antibody specific to the Flag epitope.

According to the AlphaFold model (https://alphafold.ebi.ac.uk), the Slt2 protein’s kinase domain exhibits a canonical protein kinase domain with a smaller N-terminal lobe (N-lobe) and larger C-terminal lobe (C-lobe) (**Fig 5C**), while the adjacent helix-αL adopts an elongated conformation with its N-terminal end bound to the helix-αC of N-lobe. To identify the minimum functional domain of Slt2 as well as to assess the importance of the elongated helix-αL within Slt2 kinase, we systematically removed the C-terminal residues. Subsequently, we expressed these truncated proteins (comprising of residues 1-440, 1-400 and 1-355) in the *slt2Δ* strain using a *GAL1* promoter inducible by galactose. The truncated Slt2 proteins (Slt2^1-440^ and Slt2^1-400^) consisting of the kinase domain and the helix-αL fully complemented the Slt2 function (**Fig 5D**, rows 3 and 4). However, the truncated Slt2 protein consisting of the kinase domain and a shorter helix-αL (mutant Slt2^1-355^) failed to restore the Slt2 function (**Fig 5D**, row 5). Western blot showed that the expression of truncated protein consisting of residues 1 to 355 (Slt2^1-355^) was like the WT Slt2 protein (**Fig 5E**, rows 3-6). These results indicate that C-terminal end of Slt2 protein has a redundant function. Additionally, these results suggest that the sole Slt2 kinase domain is non-functional, while the helix-αL plays an important role in its function, potentially contributing to the overall architecture of the Slt2 protein.

### 4. Both human MAP kinases ERK1 and ERK2 complement the Slt2 function under a condition of ER stress

Human ERK5 belongs to a family of CMGC kinase group(51), containing a kinase domain at its N-terminus (52). The ERK5-KD has been shown to complement the Slt2 null strain when exposed to genotoxic stresses such as HU (hydroxy urea), MMS (methyl methane sulphonate), and UV radiation(19). These findings suggest that Slt2 is the human ortholog of ERK5 (19). Consistent with this observation, we found that the expression of *Homo sapiens* ERK5-KD (residues 1-370, hsERK5) under the control of a *GAL1* promoter complemented the Slt2 function when the *slt2Δ* stain was grown on the medium containing caffeine, which is a cell wall stress-inducing agent (**Fig 6B**, row 5). However, ERK5-KD did not complement the Slt2 function when the *slt2Δ* stain was grown on the medium containing tunicamycin (**Fig 6B**, middle panel, row 5), suggesting that human ERK5 was not functional ortholog of Slt2 regarding the ER stress response.

**Figure 6:**
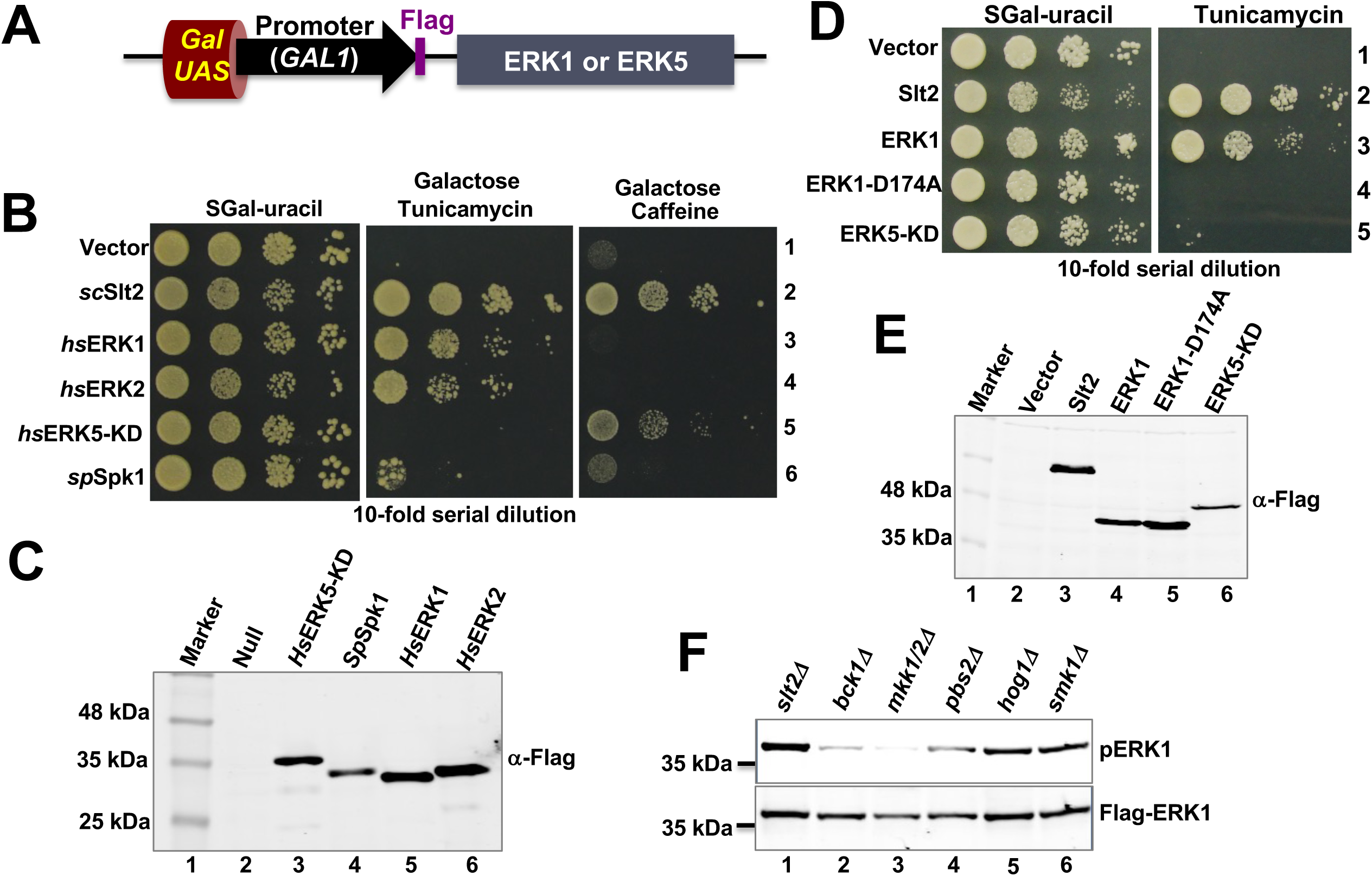
Complementation of Slt2 null strain by human MAP kinases ERK1 and ERK2. **(A)** The schematic diagram illustrates the plasmid construct for the expression ERK1 and ERK2 under the regulation of *GAL1* promoter induced by upstream activator sequence (UAS). **(B)** & (**D**) The *slt2Δ* strain containing a vector plasmid or the same vector expressing the indicated Flag-tagged MAP kinases were tested for growth on the complete synthetic (SC) medium with galactose without uracil and the same medium containing tunicamycin. **(C)** & (**E**) WCEs were prepared from yeast strain indicated in the panel (**B** and **D**) and subjected to Western blot analysis using antibody specific to the Flag epitope. **(F)** WCEs were prepared from the indicated yeast strains expressing the Flag-tagged human ERK1 and subjected to Western blot analysis using antibody specific to the phosphorylated ERK1 and ERK1.

To assess the functional ortholog of Slt2 regarding the ER stress response, we investigated whether MAP kinase ERK1, ERK2, ERK5 or p38a could compensate the loss of Slt2 kinase. A *slt2Δ* stain was transformed with a plasmid bearing the coding sequences of the *Homo sapiens* ERK1-KD (residues 42-379, *hs*ERK1), ERK2 (residues 1-360, *hs*ERK2) and p38a (residues 1-367, *hs*P38a) under a *GAL1* promoter inducible by galactose. Additionally, the *slt2Δ* stain was transformed with a plasmid bearing the coding sequences of Spk1, a MAPK kinase of the fission yeast *Schizosaccharomyces pombe (sp*Spk1)(53). Transformants were then tested for growth on the medium containing galactose (2%) with or without tunicamycin or caffeine. The *slt2Δ* stain expressing the ERK1 or ERK2 grew on the medium containing tunicamycin (0.25 μg/ml), but not on the medium containing caffeine (**Fig 6B**, rows 3 and 4, **Fig 6D**, row 3 and **Fig 6C** and **Fig 6E**). The *slt2Δ* stain containing the *sp*Spk1, the kinase inactive ERK1-D174A mutant (D = Aspartate of the catalytic HRD motif), or *hs*P38a (data not shown) did not grow on the medium containing tunicamycin (**Fig 6B**, row 6, and **Fig 6D**, row 4). Western blot shows the expression both *sp*Spk1 and ERK1-D174A proteins (**Fig 6C**, lane 4, **Fig 6E**, lane 5), suggesting that MAPK kinase of the fission yeast was unable to complement the loss of Slt2 in the budding yeast. Additionally, these results suggest that kinase activity was important for ERK1 function and yeast growth under an ER stress condition. Together, these results suggest that both ERK1 and ERK2 are functional orthologs of Slt2 kinase regarding the ER stress response.

To investigate the potential connection of upstream kinases Bck1 and Mkk1/2 with the activation of EKR1, we examined the phosphorylation status of ERK1 within its activation loop in the yeast strain lacking Bck1 (*bck1Δ*) or both Mkk1 and Mkk1 (*mkk1/2Δ*). We observed that the phosphorylation of ERK1 was substantially reduced in both *bck1Δ* and *mkk1/2Δ* strains (**Fig 6F**, lanes 2, and 3), but not in strain lacking *pbs1Δ*, *hog1Δ* and *smk1Δ* strains (**Fig 6F**, lanes 4, 5, and 6). These results suggest that Bck1 and Mkk1/2 activates ERK1, akin to their role in activation of the kinase Slt2.

### 5. Activation of MAP kinase ERK1 during ER stress in primary human cells

Using a human embryonic kidney (HEK-293) cell line, we examined how ERK1 was phosphorylated under a condition of ER stress. Cells were incubated with the ER stressor tunicamycin (5μg/ml) for 2 and 4 hours as described in the **Materials and Methods**. Whole cell extracts were prepared and subjected to Western blot analysis. As anticipated, expression of the ER stress marker BIP was increased with upon exposure of cell to tunicamycin (**Fig 7A** and **7B**). The basal level of ERK1/2 phosphorylation was higher in the immortalized HEK293 cells (see double protein bands in **Figs 7A** and **7B**), indicating that ERK1/2 signaling pathway might play a critical role in cell proliferation. No significant increase in the level of ERK phosphorylation was observed when cells were exposed to tunicamycin (**Fig 7A** and **7B**). Additionally, we examined the level of ERK phosphorylation in serum-deprived HEK-293 cells in the presence of tunicamycin. No detectable increase in the level of ERK phosphorylation was observed (data not shown). Together, we assumed that immortal HEK293 cells might not be a right *in vitro* tool for studying the ERK1/2 phosphorylation during the ER stress. Therefore, we examined phosphorylation of ERK1 in the primary cells (i.e., ovarian surface epithelial cells) grown in the presence and absence of tunicamycin.

**Figure 7:**
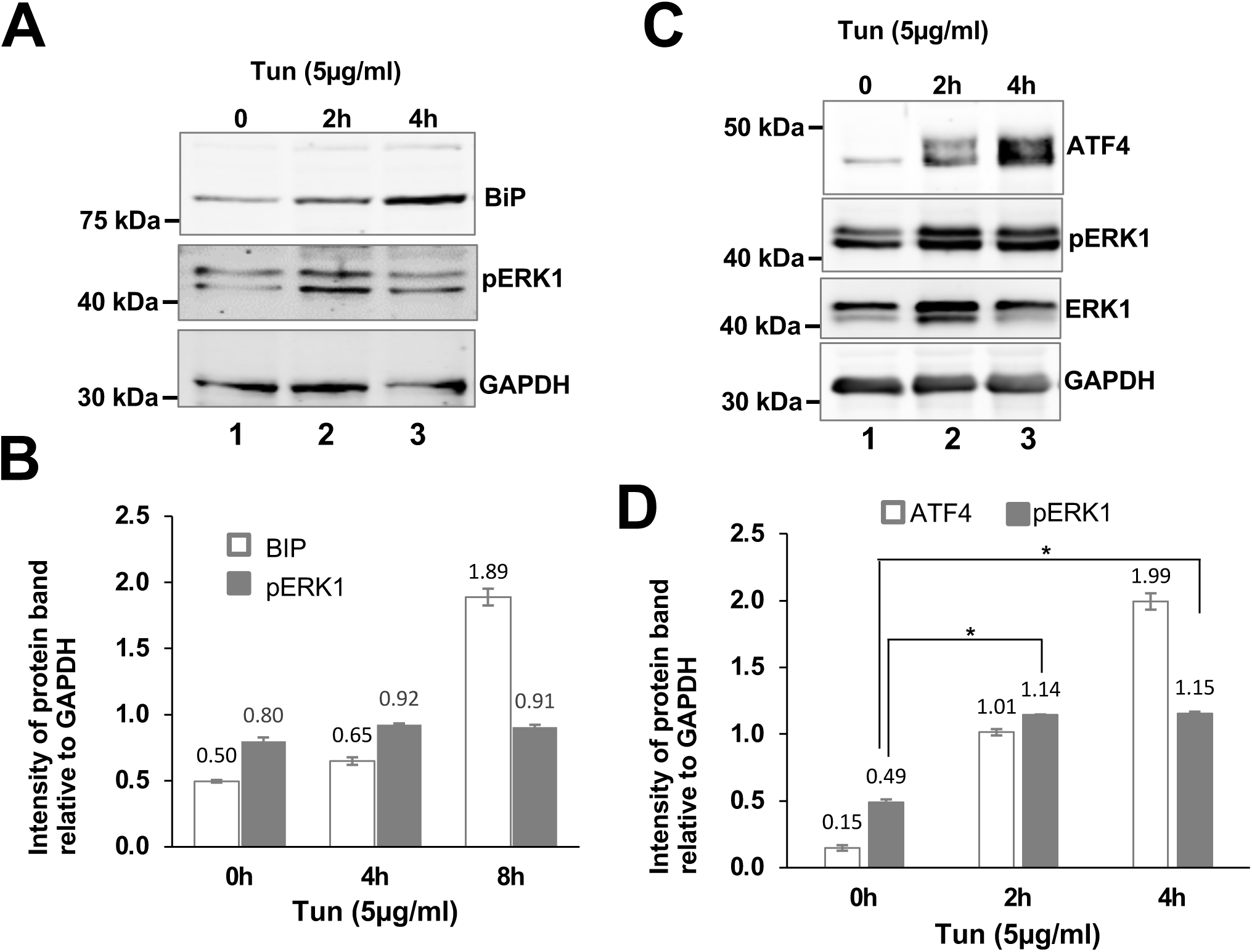
Increased level of phospho-ERK1 in Ovarian Surface Epithelial (OSE) cells grown in the presence of ER stressor tunicamycin. **(A)** WCEs were prepared from the HEK293 cells grown in the presence of tunicamycin and subjected to Western blot analysis using antibodies specific to the BiP, phosphorylated ERK1 and GAPDH. The intensities of protein bands were quantified using the ImageJ software. **(B)** The bar diagram depicts the relative intensities of BiP and pERK1 protein bands. **(C)** WCEs were prepared from the Ovarian Surface Epithelial (OSE) cells grown in the presence of tunicamycin and subjected to Western blot analysis using antibodies specific to the phosphorylatedERK1, ERK1, ATF4, and GAPDH. The intensities of protein bands were quantified using the ImageJ software. (Lower panel) **(D)** The bar diagram depicts the relative intensities of ATF4 and pERK1 protein bands.

We collected samples of normal ovarian epithelial tissues from individuals with the tumor-free ovary, unilateral ovarian cancer or patients with other gynecological cancers not involving the ovary, following the Institutional Review Board approved protocol at Medical College of Wisconsin. Ovarian surface epithelial cells (OSE) were cultured by scraping the surface epithelium of normal ovarian tissues. OSE cells were cultured in MCDB105/M199 medium (Sigma-Aldrich, St. Louis, MO) (1:1 mixture) supplemented with 10% FBS, human epidermal growth factor and insulin in the presence of tunicamycin. As expected, the expression of the ER stress marker ATF4 was increased when exposed of cells to tunicamycin (**Fig 7C** and **7D**). We observed that the primary cells exhibited a basal level of ERK1/2 phosphorylation. Tunicamycin-treated cells displayed approximately a 2-fold increase in the ERK1/2 phosphorylation (**Fig 7C** and **7D**). Taken together, these findings indicate that ER stress activates both kinases ERK1 and ERK2.

## Discussion

We demonstrate that, in the budding yeast, the cellular response to ER stress involves at least two distinct phases (**Fig 4**). The initial phase is primarily mediated by the active Ire1, resulting in translational de-repression of *HAC1* mRNA. The second phase, likely occurring at slower pace, involves the activation of Slt2. In this slower response, Ire1 pathway is likely attenuated, leading to reduced expression of Hac1 protein (**Fig 4**), while the Slt2-mediated responses come into play to promote adaptation and recovery. We also provide compelling evidence that the MAP kinase Slt2 plays a crucial role in UPR by directly influencing both splicing and translation of *HAC1* mRNA (**Fig 2**), where Slt2 serves as a key mediator in the activation of UPR genes through an alternative route. Our results also reveal that Hac1 protein alone is insufficient for adapting to ER stress (**Fig 3**). Notably, we are the first to demonstrate that, in the context of the ER stress response, human counterparts of Slt2 kinase are ERK1 and ERK2 (**Figs 6** and **7**).

### The yeast MAP kinase Slt2 pathway

The MAP kinase Slt2 is primarily known for its involvement in the cell wall integrity (CWI) signaling pathway, which is essential for maintaining the shape and integrity of cells during both vegetative growth and mating process(22,54). Activation of the CWI pathway is initiated in response to several environmental stresses such as heat shock(55), hypoosmotic stress(21) and actin depolymerization(56). For the CWI pathway, two key surface sensors are the transmembrane proteins Wsc1 and Mid2(57), which likely transmit the extracellular signals to GDP/GTP exchange factor Rom2 and the small GTP-binding protein Rho1(57). Rho1 binds and activates Pkc1, which in turn activates the terminal Slt2 MAPK module composed of Bck1 and MKK1/2 and Slt2 (58). Additionally, multiple studies highlight that Slt2 becomes active in response to ER stress (46,59,60). However, the molecular association between Slt2, CWI and Ire1 pathways, and their relationship to maintaining cellular proteostasis during ER stress, remain enigmatic. In other words, the mechanisms by which the stress caused by protein folding defects inside the ER communicates with the cytosolic MAPK Slt2 remains to be determined (**Fig 8**).

**Figure 8:**
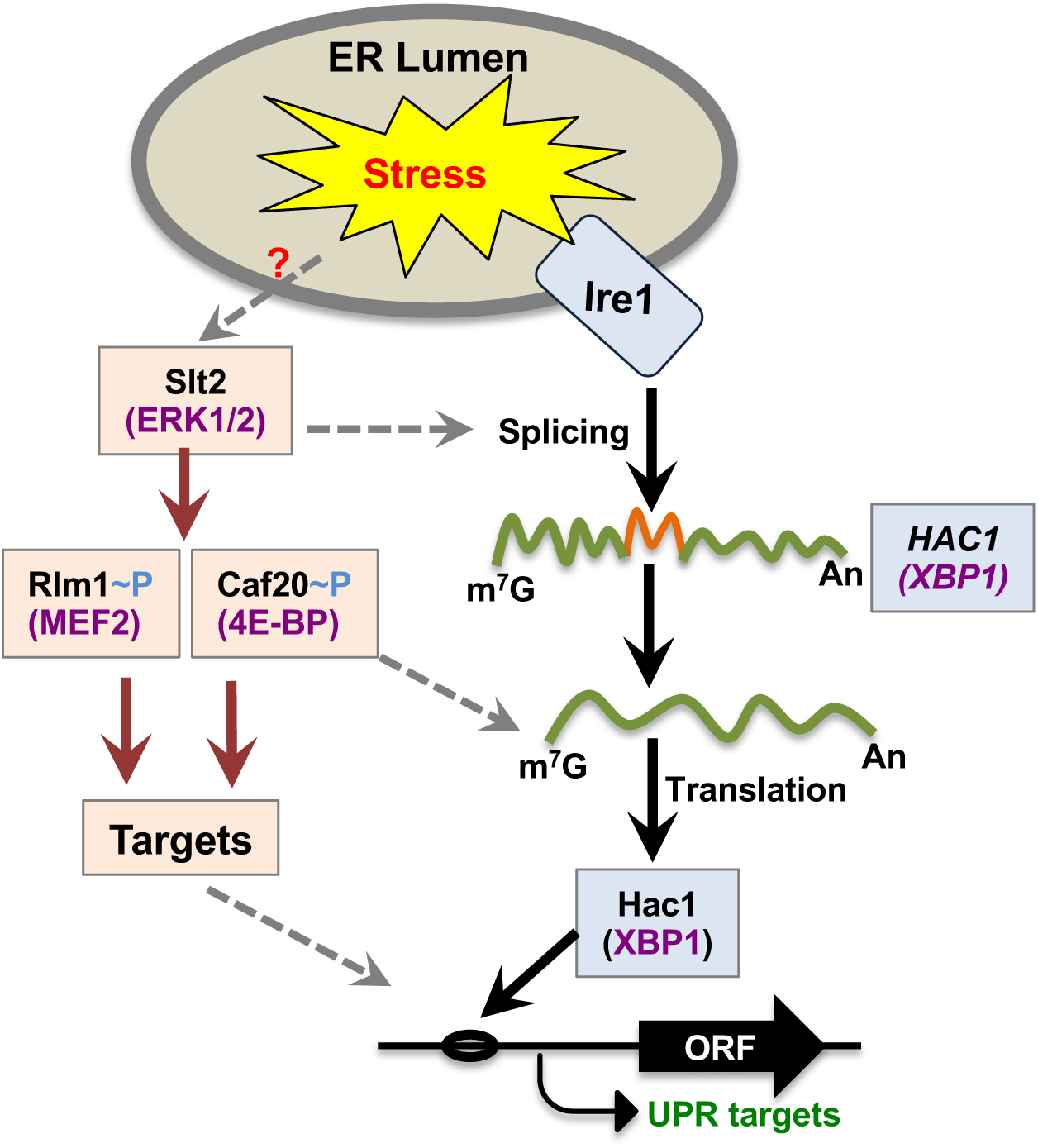
Proposed link between Ire1 and Slt2 signaling pathways during ER Stress. The ER stress activates the dual kinase and RNase Ire1 (light blue box) that removes the intronic sequence (orange wavy line) from the *HAC1* pre-mRNA (green wavy line). The matured *HAC1* mRNA then translates Hac1 protein that binds to the upstream sequence of the UPR genes, thus activating their transcription. The ER stress also activates the MAP kinase Slt2 pathway (colored in brick red) that promotes *HAC1* mRNA splicing by unknown mechanisms. The active Slt2 can phosphorylate and activate the transcriptional activator Rlm1 as well as translational suppressor Caf40, an eIF4E binding protein. Rlm1 might collaborate with Hac1 and modulate the transactivation of UPR genes. Phosphorylation of Caf20 by Slt2 might serve to inactivate its inhibitory effect on translation. The human counterparts of yeast proteins are shown in purple color.

The active kinase Slt2 is known to phosphorylate a wide variety of substrates, including transcription factors like Rlm1(61) and Swi4(62), ribosomal chaperone Ssb2(63), eIF4E-binding protein Caf20(64), transcriptional silencer Sir3(65), PKA regulatory subunit Bcy1(66), cell cycle regulator cyclin C (67), calcineurin regulator Rcn2(68), Golgi-associated adaptor Gga1(68), phosphatase Msg5(68), and the subunit of RNA polII (69). These Slt2 substrates, once phosphorylated and activated, likely play crucial roles in regulating several cellular processes, including the control of cell growth and differentiation, and the coordination of responses to various forms of stress. However, the specific mechanisms by which Slt2 phosphorylates any of these substrates, or any potentially undiscovered substrates, as well as its role in contributing to ER stress response, remain to be fully elucidated.

### The Slt2 pathway and ER stress-dependent proteostasis

The elevated levels of phosphorylation within the activation loop of Slt2 following a 2-hour exposure to ER stressor (**Fig 4**) suggest that Slt2’s involvement in ER stress response likely extends beyond its function in activating the Ire1-Hac1 signaling pathway. This notion is further supported by previous studies that the *slt2Δire1Δ* double mutant displays greater sensitivity to ER stress compared to the corresponding single mutants (46). Together, it appears that Slt2 pathway initiates an additional signaling pathway specifically in response to prolonged ER stress.

A study conducted by Rousseau and Bertolotti (2016) show that the expression levels of chaperones for the 26S proteasomal complex (e.g., Adc17, Hsm3, Nas6, Nas2 and Rpn14) were significantly reduced in the *slt2Δ* strain that was grown under a condition of ER stress induced by tunicamycin for 4 hours(48). These findings led them to propose that the tunicamycin sensitivity of the *slt2Δ* strain is likely due to defects in assembly of the functional 26S proteasome complex that clears the unfolded proteins from cells during ER stress. However, yeast strain lacking either of these chaperones grew normally on the tunicamycin medium (data not shown), underscoring the importance of Adc17, Hsm3, Nas6, Nas2 or Rpn14 in ER stress response. So, the precise role of Slt2 in ER stress remains to be conclusively determined.

In this study, we show that the *slt2Δ* strain expressing the Hac1 protein displayed impaired growth in the presence of tunicamycin (**Fig 3** and **Fig 4**). These findings suggest that the Hac1-mediated UPR relies on signals from multiple sources, including the Slt2 pathway. These signals are presumed to generate specific transcriptional program appropriate for the specific physiological requirements of cells. One potential scenario involves a specific signaling input originating from the Slt2 pathway, achieved through the phosphorylation of one of its reported substrates. This phosphorylated substrate may collaborate with the Hac1 to activate a particular physiological response.

Research on the involvement of Slt2 in UPR(46,60,70) have primarily focused on the *slt2Δ* cell grown in the presence of ER stressors for a short duration (one hour). For instance, Chen at al. (2005) show comparable levels of *HAC1* mRNA splicing between WT and *slt2Δ* strains after one hour of growth under an ER stress condition(46). Similar findings were obtained in our experiments (data not shown). However, these results alone do not explain why the *slt2Δ* strain is severely sensitivity to tunicamycin (**Fig 1B**). One plausible explanation is that only one of hour of stress is insufficient to see any effect because the activation of pro-survival strategies relies upon the duration of stress signals. Thus, we extended the ER stress in the *slt2Δ* strain for at least 2 hours and monitored the *HAC1* mRNA splicing and translation. The notable reductions (>50% compared to the isogenic WT cells) in both splicing and translation of *HAC1* mRNA were observed (**Fig 2**), This decrease in splicing may be associated with a decrease in the functionality of the Ire1-RNase and/or tRNA ligase functions, while the decrease in the Hac1 protein expression is associated with the reduced translation efficiency of the spliced *HAC1* mRNA (**Fig 8**). Collectively, our data strongly suggest that Slt2 functions in a signaling pathway that directly connects the Ire1-Hac1 mediated UPR.

In ongoing studies, we are investigating if the translational efficiency of *HAC1* mRNA and Hac1’s transactivation of UPR genes could be influenced by two Slt2 substrates Rlm1 (an ortholog of human MEF2) and Caf20 (an ortholog of human eIF4E-BP) (**Fig 8**). Rlm1 is a MADS-box transcription factor that binds to a conserved 5’-CA(A/T)_6_G-3’ DNA element at the promoter(71), leading to activation of several genes in response to cell wall stress(61). Notably, yeast cells lacking Rlm1 display partial sensitivity to tunicamycin (data not shown), suggesting that Rlm1 plays a role in response to ER stress. Given these observations, it is plausible that Rlm1 might collaborate with Hac1 and modulate the transactivation of UPR genes in the context of sustainable UPR (**Fig 8**). In contrast to Rlm1, the eIF4E-binding protein Caf20 likely sequesters the cap binding protein eIF4E, thereby inhibiting the translation efficiency (72,73). It has also been speculated that Caf20 binds to the 3’-UTR of many mRNAs(74), thus inhibiting translation by interfering with the interaction between eIF4G and PABP (<u>p</u>oly<u>A</u>-<u>b</u>inding <u>p</u>rotein) that enhances the translation efficiency by circularizing mRNA(74). Therefore, it is possible that Caf20 could potentially interact with and capture eIF4E, leading to a reduction in the eIF4E•eIF4G interaction, the formation of eIF4F, and ultimately resulting in reduced translation efficiency of spliced *HAC1* mRNA. Phosphorylation of Caf20 by Slt2 and other kinase, such as Psk2(75), might serve to inactivate its inhibitory effect (**Fig 8**).

### The Helix-**α**L in MAP kinase function

The helix-αC is a dynamic regulatory element within the protein kinase domain(76). In the active state of a kinase domain, the helix-αC’s N-terminal end engages with the phosphorylated activation loop. The distance between this N-terminal end of the helix-αC and the phosphorylated activation loop defines the open and closed conformations of the kinase domain which is crucial for the kinase catalysis(76). The helix-αC of ERK1/2 kinase domain interacts with the helix-αL located at the adjoining C-terminal end(50). In this study, we show that the analogous helix-αL of the Slt2 kinase domain is important for its role in ER stress response (**Fig 5B** and **Fig 5C**). These findings indicate that the helix-αC•helix-αL interaction likely controls the overall conformation and activation of both kinases ERK1 and ERK2. However, the precise physiological evidence supporting this functional interaction and elucidating the activation mechanisms, particularly the involvement of extended helix-αL in Slt2 kinase, remains to be determined.

### The MAPK substrates in UPR

The MAP kinase Slt2 displays its closest sequence similarity with members of the mammalian MAPK family, including ERK1, ERK2 and ERK5 (**Fig 6B**). It is worth mentioning that none of these mammalian MAP kinases displayed more than 50% identity to Slt2. Nonetheless, the expression of either ERK1 or ERK2 could compensate the loss of Slt2 kinase in the context of ER stress response (**Fig 6B** and **Fig 6C**), whereas ERK5, despite being regarded as a functional ortholog of yeast Slt2 (19), did not exhibit this same compensatory effect. To be noted that the expression of the kinase domain of ERK5 could compensate the loss of Slt2 kinase in the context of genotoxic stresses (19). Furthermore, we observe that fission yeast Spk1, an ortholog of Slt2, did not have any compensatory effect regarding the cellular response to ER stress (**Fig 6B**). These observations collectively suggest that significant differences exist in the regulation of MAPK pathways across different species, despite the evolutionarily conservation of these stress-related pathways.

ERK1 and ERK2 appear to be constitutively active in cultured human cells (**Fig 7A**). This could be attributed the fact that the active ERK1/2 are known to phosphorylate a diverse group of substrates (e.g., protein kinases, phosphatases, and regulators of apoptosis) to control multiple signaling pathways(77–80). However, the mechanisms how ERK1/2 can phosphorylate hundreds of substrates(81) remain unknown, although it has been shown that ERK1/2 specificity depends their location (e.g., nucleus and cytoplasm), the specific needs (e.g., stress response) and the substrate docking site (e.g., DEF motif(82)). While ERK1 is known to phosphorylate ∼600 known substrates(77,81), the real challenge is to identify a UPR-specific substrate.

### Experimental procedures

#### Yeast strains, growth, and gene disruption

Yeast *S. cerevisiae* strains were grown in the standard medium (1% yeast extract, 2% peptone and 2% dextrose [YEPD]) or defined synthetic complete (SC, 0.17% yeast nitrogen base [YNB], 0.5% ammonium sulphate, 2% glucose and all amino acids). The genomic DNA of the *ire1::HphMX* strain was used as a template to amplify the *HphMX* cassette using primers annealing ∼200-bases upstream and downstream of the *IRE1* open reading frame. The amplified PCR product was used to disrupt the *IRE1* gene of the *slt2::KanMX* strain. The list of yeast strains used in this study is shown in **Table 1**.

**Table 1.** List of yeast strains used in this study.

| Strain Name | Genotype | Reference |
| --- | --- | --- |
| WT (BY4741) | *MATa his3-Δ1 leu2-Δ0 met5-Δ0 ura3-Δ0* | Deletion collection |
| *ire1Δ* | *MATa his3-Δ1 leu2-Δ0 met5-Δ0 ura3-Δ0 ire1::KanMX* | Deletion collection |
| *ire1Δ* | *MATa his3-Δ1 leu2-Δ0 met5-Δ0 ura3-Δ0 ire1::HphMx* | This study |
| *ste20Δ* | *MATa his3-Δ1 leu2-Δ0 met5-Δ0 ura3-Δ0 ste20::KanMX* | Deletion collection |
| *ste1Δ* | *MATa his3-Δ1 leu2-Δ0 met5-Δ0 ura3-Δ0 ste1::KanMX* | Deletion collection |
| *ste7Δ* | *MATa his3-Δ1 leu2-Δ0 met5-Δ0 ura3-Δ0 ste7::KanMX* | Deletion collection |
| *fus3Δ* | *MATa his3-Δ1 leu2-Δ0 met5-Δ0 ura3-Δ0 fus3::KanMX* | Deletion collection |
| *kss1Δ* | *MATa his3-Δ1 leu2-Δ0 met5-Δ0 ura3-Δ0 kss1::KanMX* | Deletion collection |
| *pbs2Δ* | *MATa his3-Δ1 leu2-Δ0 met5-Δ0 ura3-Δ0 pbs2::KanMX* | Deletion collection |
| *hog1Δ* | *MATa his3-Δ1 leu2-Δ0 met5-Δ0 ura3-Δ0 hog1::KanMX* | Deletion collection |
| *ssk1Δ* | *MATa his3-Δ1 leu2-Δ0 met5-Δ0 ura3-Δ0 ssk1::KanMX* | Deletion collection |
| *ssk2Δ* | *MATa his3-Δ1 leu2-Δ0 met5-Δ0 ura3-Δ0 ssk2::KanMX* | Deletion collection |
| *ssk22Δ* | *MATa his3-Δ1 leu2-Δ0 met5-Δ0 ura3-Δ0*<br>*ssk22::KanMX* | Deletion collection |
| *bck1Δ* | *MATa his3-Δ1 leu2-Δ0 met5-Δ0 ura3-Δ0 bck1::KanMX* | Deletion collection |
| *mkk1Δ, mkk2Δ*<br>*(mkk1/2Δ)* | *MATa his3-Δ1 leu2-Δ0 met5-Δ0 ura3-Δ0 mkk1::KanMX*<br>*mkk2::NatMX* | Deletion collection |
| *slt2Δ* | *MATa his3-Δ1 leu2-Δ0 met5-Δ0 ura3-Δ0 slt2::KanMX* | Deletion collection |
| *smk1Δ* | *MATa his3-Δ1 leu2-Δ0 met5-Δ0 ura3-Δ0 smk1::KanMX* | Deletion collection |
| *ire1Δslt2Δ* | *MATa his3-Δ1 leu2-Δ0 met5-Δ0 ura3-Δ0 slt2::KanMX*<br>*ire1::HphMX* | This study |

#### Preparation of plasmids

Plasmids were generated using the standard gene manipulation techniques. Mutation was generated by fusion PCR using standard protocols. The desired mutation in each plasmid was confirmed by Sanger sequencing. The list of plasmids used in this study are shown in **Table 2**.

**Table 2:** List of plasmids used in this study.

| Plasmid Name | Plasmid description | Reference or study |
| --- | --- | --- |
| D4 | pRS316, low-copy-numbered URA3 vector | Lab collection |
| D1984 | pYES2 vector | Lab collection |
| D69 | HAC1<sup>c</sup> (intron less) in D4 | Uppala JK et al., 2021 |
| D63 | HAC1-WT in D4 | Sathe L et al., 2015 |
| D1694 | HAC1-G771A in D4 | Sathe L et al., 2015 |
| D2305 | FLAG-SLT2 in D1984 | This study |
| D2716 | FLAG-SLT2<sup>1-440</sup> in D1984 | This study |
| D2720 | FLAG-SLT2<sup>1-400</sup> in D1984 | This study |
| D2313 | FLAG-SLT2<sup>1-355</sup> in D1984 | This study |
| D2276 | Human FLAG-ERK1 in D1984 | This study |
| D2277 | Human FLAG-ERK2 in D1984 | This study |
| D2255 | Human FLAG-ERK5-KD in D1984 | This study |
| D2197 | Schizosaccharomyces pombe SPK1 in D1984 | This study |
| D2312 | Human FLAG-ERK1-D174A in D1984 | This study |
| D2480 | Human FLAG-P38α in D1984 | This study |

#### Whole cell extract preparation and Western blot analysis

Yeast cells were grown in YEPD or Synthetic complete (SC) medium without appropriate nutrients until the OD_600_ value reached ∼0.5 to 0.6. DTT (5 mM) or tunicamycin (0.5 μg/ml) was added to the medium to induce ER stress, and cells were harvested after 2 hours (unless otherwise indicated). Whole cell extracts (WCEs) were prepared by TCA method as described previously (83). Proteins were then fractioned by SDS–PAGE and Western blot analysis was performed using appropriate antibody (See **Table 3**). All experiments were repeated at least two times.

**Table 3:** List of antibodies used in this study.

| Antibody | Source | Catalog number | Dilution |
| --- | --- | --- | --- |
| Anti-BiP | Santa Cruz<br>Biotechnology | SC-376768 | 1: 1000 |
| Anti-ATF4 | Cell signaling | 11815S | 1: 1000 |
| Anti-pERK1 | Cell signaling | D11A8 | 1: 1000 |
| Anti-ERK1/2 | Cell signaling | 9102 | 1: 2000 |
| Anti-GAPDH | Santa Cruz<br>Biotechnology | SC-32233 | 1: 1000 |
| Anti-Flag | Sigma-Aldrich | F3165 | 1: 1000 |
| Anti-pSlt2 | Cell signaling | 9101S | 1: 1000 |
| Anti-Slt2 | Santa Cruz<br>Biotechnology | SC-133189 | 1: 1000 |
| Anti- Pgk1 | Invitrogen | 459250 | 1:1000 |
| Anti-Hac1 | Dey lab | - | 1: 500 |

#### RNA analysis and reverse transcriptase (RT)-PCR

Yeast cells were grown in YEPD or SC medium without appropriate nutrients at 30°C until they reached an OD_600_ value between 0.5 and 0.6. The ER stressor DTT (5 mM) or tunicamycin (1 μg/ml) was added to the medium and cells were grown further for another 2 hours (unless otherwise indicated). Cells were harvested and total RNA was isolated using the RNeasy mini kit (Qiagen). Purified RNA was quantified using a Nanadrop spectrophotometer (ND-1000, Thermo Scientific) and used to synthesize the first strand cDNA by a Superscript^TM^-III reverse transcriptase (Invitrogen 18080-093) and a reverse primer (5-CCCACCAACAGCGATAATAACGAG-3) that corresponded to nucleotides +1002 to 1025. To assay *HAC1* mRNA splicing, the synthetic cDNA was then PCR-amplified using a forward primer (5-CGCAATCGAACTTGGCTATCCCTA CC-3) that corresponded to nucleotides +35 to 60 and a reverse primer (5-CCCACCA ACAGCGATAATAACGAG-3) that corresponded to nucleotides +1002 to 1025. The PCR-amplified products were then run on a 1.5% agarose gel to separate spliced (HAC1^s^) and un-spliced (HAC1^u^) forms of *HAC1* mRNA. Quantities of HAC1^s^ and HAC1^u^ were measured using ImageJ. Experiments were repeated at least two times.

#### Human cell culturing and whole cell extract (WCE) preparation

HEK293T cells were cultured in a high glucose Dulbecco’s modified Eagle’s medium (DMEM) medium supplemented with 10% fetal bovine serum, 100 U of penicillin G, 100 µg/ml streptomycin, and 6 mM L-glutamine. Cells were maintained at 37°C with 5% CO_2_. Cells were seeded in a 6-well cell culture plate one day before the drug treatment. Cells were treated with tunicamycin (5 µg/ml) for 2 and 4 hours and washed with 1X phosphate buffered saline (PBS). Cells were then incubated in a lysis buffer (20 mM Tris-HCl (pH 8.0), 80 mM KCl, 1 mM EDTA, 0.5% NP-40, 1 mM DTT, 1X protease inhibitor cocktail and 1X phosphatase inhibitor cocktail) for 10 min and lysed by pipetting followed by vortex for 15 min at 4°C. Cell extract was then centrifuged at 12,000 g for 15 min at 4°C. The clear supernatant containing the whole cell extracts were collected and subjected to Western blot analysis using appropriate antibody.

#### Analysis of biomolecule structures

The coordinates of the predicted Slt2 structure were retrieved from the AlphaFold web site (https://alphafold.ebi.ac.uk) and analyzed using the free PyMol software (https://www.pymol.org/pymol)

#### Data analysis and densitometry analysis

The densitometry analysis was performed using the NIH ImageJ gel analysis software(84). All bands at the correct molecular weight ± approximately 5 kDa were analyzed thrice as the signal for that target protein. The average of three measurements with SD (standard deviation) are shown in each plot. Comparisons are made against the same sample lysate. Each Western blot was repeated at least twice.

## Acknowledgements

We would like to thank Prof. David Frick for reading and editing the manuscript. This work was supported by grants to M.D. from the U.S. National Institutes of Health (R01 GM124183-04) and UWM graduate school (Discovery and Innovation Grant).

## References

1. Keshet, Y., and Seger, R. (2010) The MAP kinase signaling cascades: a system of hundreds of components regulates a diverse array of physiological functions. Methods Mol Biol 661, 3–38

2. Morrison, D. K. (2012) MAP kinase pathways. Cold Spring Harb Perspect Biol 4

3. Qi, M., and Elion, E. A. (2005) MAP kinase pathways. J Cell Sci 118, 3569–3572

4. Boulton, T. G., Nye, S. H., Robbins, D. J., Ip, N. Y., Radziejewska, E., Morgenbesser, S. D., DePinho, R. A., Panayotatos, N., Cobb, M. H., and Yancopoulos, G. D. (1991) ERKs: a family of protein-serine/threonine kinases that are activated and tyrosine phosphorylated in response to insulin and NGF. Cell 65, 663–675

5. Boulton, T. G., Yancopoulos, G. D., Gregory, J. S., Slaughter, C., Moomaw, C., Hsu, J., and Cobb, M. H. (1990) An insulin-stimulated protein kinase similar to yeast kinases involved in cell cycle control. Science 249, 64–67

6. English, J. M., Vanderbilt, C. A., Xu, S., Marcus, S., and Cobb, M. H. (1995) Isolation of MEK5 and differential expression of alternatively spliced forms. J Biol Chem 270, 28897–28902

7. Derijard, B., Hibi, M., Wu, I. H., Barrett, T., Su, B., Deng, T., Karin, M., and Davis, R. J. (1994) JNK1: a protein kinase stimulated by UV light and Ha-Ras that binds and phosphorylates the c-Jun activation domain. Cell 76, 1025–1037

8. Hibi, M., Lin, A., Smeal, T., Minden, A., and Karin, M. (1993) Identification of an oncoprotein- and UV-responsive protein kinase that binds and potentiates the c-Jun activation domain. Genes Dev 7, 2135–2148

9. Kyriakis, J. M., Banerjee, P., Nikolakaki, E., Dai, T., Rubie, E. A., Ahmad, M. F., Avruch, J., and Woodgett, J. R. (1994) The stress-activated protein kinase subfamily of c-Jun kinases. Nature 369, 156–160

10. Kassouf, T., and Sumara, G. (2020) Impact of Conventional and Atypical MAPKs on the Development of Metabolic Diseases. Biomolecules 10

11. Han, J., Lee, J. D., Bibbs, L., and Ulevitch, R. J. (1994) A MAP kinase targeted by endotoxin and hyperosmolarity in mammalian cells. Science 265, 808–811

12. Lee, J. C., Laydon, J. T., McDonnell, P. C., Gallagher, T. F., Kumar, S., Green, D., McNulty, D., Blumenthal, M. J., Heys, J. R., Landvatter, S. W., Strickler, J. E., McLaughlin, M. M., Siemens, I. R., Fisher, S. M., Livi, G. P., White, J. R., Adams, J. L., and Young, P. R. (1994) A protein kinase involved in the regulation of inflammatory cytokine biosynthesis. Nature 372, 739–746

13. Kato, Y., Kravchenko, V. V., Tapping, R. I., Han, J., Ulevitch, R. J., and Lee, J. D. (1997) BMK1/ERK5 regulates serum-induced early gene expression through transcription factor MEF2C. EMBO J 16, 7054–7066

14. Tibbles, L. A., and Woodgett, J. R. (1999) The stress-activated protein kinase pathways. Cell Mol Life Sci 55, 1230–1254

15. Wagner, E. F., and Nebreda, A. R. (2009) Signal integration by JNK and p38 MAPK pathways in cancer development. Nat Rev Cancer 9, 537–549

16. Canovas, B., and Nebreda, A. R. (2021) Diversity and versatility of p38 kinase signalling in health and disease. Nat Rev Mol Cell Biol 22, 346–366

17. Gustin, M. C., Albertyn, J., Alexander, M., and Davenport, K. (1998) MAP kinase pathways in the yeast Saccharomyces cerevisiae. Microbiol Mol Biol Rev 62, 1264–1300

18. Brewster, J. L., de Valoir, T., Dwyer, N. D., Winter, E., and Gustin, M. C. (1993) An osmosensing signal transduction pathway in yeast. Science 259, 1760–1763

19. Truman, A. W., Millson, S. H., Nuttall, J. M., King, V., Mollapour, M., Prodromou, C., Pearl, L. H., and Piper, P. W. (2006) Expressed in the yeast Saccharomyces cerevisiae, human ERK5 is a client of the Hsp90 chaperone that complements loss of the Slt2p (Mpk1p) cell integrity stress-activated protein kinase. Eukaryot Cell 5, 1914–1924

20. Kamada, Y., Jung, U. S., Piotrowski, J., and Levin, D. E. (1995) The protein kinase C-activated MAP kinase pathway of Saccharomyces cerevisiae mediates a novel aspect of the heat shock response. Genes Dev 9, 1559–1571

21. Davenport, K. R., Sohaskey, M., Kamada, Y., Levin, D. E., and Gustin, M. C. (1995) A second osmosensing signal transduction pathway in yeast. Hypotonic shock activates the PKC1 protein kinase-regulated cell integrity pathway. J Biol Chem 270, 30157–30161

22. Zarzov, P., Mazzoni, C., and Mann, C. (1996) The SLT2(MPK1) MAP kinase is activated during periods of polarized cell growth in yeast. EMBO J 15, 83–91

23. Irie, K., Takase, M., Lee, K. S., Levin, D. E., Araki, H., Matsumoto, K., and Oshima, Y. (1993) MKK1 and MKK2, which encode Saccharomyces cerevisiae mitogen-activated protein kinase-kinase homologs, function in the pathway mediated by protein kinase C. Mol Cell Biol 13, 3076–3083

24. Levin, D. E. (2011) Regulation of cell wall biogenesis in Saccharomyces cerevisiae: the cell wall integrity signaling pathway. Genetics 189, 1145–1175

25. Darling, N. J., and Cook, S. J. (2014) The role of MAPK signalling pathways in the response to endoplasmic reticulum stress. Biochim Biophys Acta 1843, 2150–2163

26. Wang, X. Z., Harding, H. P., Zhang, Y., Jolicoeur, E. M., Kuroda, M., and Ron, D. (1998) Cloning of mammalian Ire1 reveals diversity in the ER stress responses. EMBO J 17, 5708–5717

27. Tirasophon, W., Welihinda, A. A., and Kaufman, R. J. (1998) A stress response pathway from the endoplasmic reticulum to the nucleus requires a novel bifunctional protein kinase/endoribonuclease (Ire1p) in mammalian cells. Genes Dev 12, 1812–1824

28. Sidrauski, C., and Walter, P. (1997) The transmembrane kinase Ire1p is a site-specific endonuclease that initiates mRNA splicing in the unfolded protein response. Cell 90, 1031–1039

29. Harding, H. P., Zhang, Y., and Ron, D. (1999) Protein translation and folding are coupled by an endoplasmic-reticulum-resident kinase. Nature 397, 271–274

30. Haze, K., Yoshida, H., Yanagi, H., Yura, T., and Mori, K. (1999) Mammalian transcription factor ATF6 is synthesized as a transmembrane protein and activated by proteolysis in response to endoplasmic reticulum stress. Mol Biol Cell 10, 3787–3799

31. Schroder, M. (2008) Endoplasmic reticulum stress responses. Cell Mol Life Sci 65, 862–894

32. Walter, P., and Ron, D. (2011) The unfolded protein response: from stress pathway to homeostatic regulation. Science 334, 1081–1086

33. Ron, D., and Walter, P. (2007) Signal integration in the endoplasmic reticulum unfolded protein response. Nat Rev Mol Cell Bio 8, 519–529

34. Ruggiano, A., Foresti, O., and Carvalho, P. (2014) Quality control: ER-associated degradation: protein quality control and beyond. J Cell Biol 204, 869–879

35. Olzmann, J. A., Kopito, R. R., and Christianson, J. C. (2013) The mammalian endoplasmic reticulum-associated degradation system. Cold Spring Harb Perspect Biol 5

36. Brodsky, J. L. (2012) Cleaning up: ER-associated degradation to the rescue. Cell 151, 1163–1167

37. Sidrauski, C., Cox, J. S., and Walter, P. (1996) tRNA ligase is required for regulated mRNA splicing in the unfolded protein response. Cell 87, 405–413

38. Lu, Y., Liang, F. X., and Wang, X. (2014) A synthetic biology approach identifies the mammalian UPR RNA ligase RtcB. Mol Cell 55, 758–770

39. Vattem, K. M., and Wek, R. C. (2004) Reinitiation involving upstream ORFs regulates ATF4 mRNA translation in mammalian cells. Proc Natl Acad Sci U S A 101, 11269–11274

40. Hollien, J., and Weissman, J. S. (2006) Decay of endoplasmic reticulum-localized mRNAs during the unfolded protein response. Science 313, 104–107

41. Hollien, J., Lin, J. H., Li, H., Stevens, N., Walter, P., and Weissman, J. S. (2009) Regulated Ire1-dependent decay of messenger RNAs in mammalian cells. J Cell Biol 186, 323–331

42. Chang, T. K., Lawrence, D. A., Lu, M., Tan, J., Harnoss, J. M., Marsters, S. A., Liu, P., Sandoval, W., Martin, S. E., and Ashkenazi, A. (2018) Coordination between Two Branches of the Unfolded Protein Response Determines Apoptotic Cell Fate. Mol Cell 71, 629–636 e625

43. Morishima, N., Nakanishi, K., and Nakano, A. (2011) Activating transcription factor-6 (ATF6) mediates apoptosis with reduction of myeloid cell leukemia sequence 1 (Mcl-1) protein via induction of WW domain binding protein 1. J Biol Chem 286, 35227–35235

44. Baetz, K., Moffat, J., Haynes, J., Chang, M., and Andrews, B. (2001) Transcriptional coregulation by the cell integrity mitogen-activated protein kinase Slt2 and the cell cycle regulator Swi4. Mol Cell Biol 21, 6515–6528

45. Watanabe, Y., Irie, K., and Matsumoto, K. (1995) Yeast RLM1 encodes a serum response factor-like protein that may function downstream of the Mpk1 (Slt2) mitogen-activated protein kinase pathway. Mol Cell Biol 15, 5740–5749

46. Chen, Y., Feldman, D. E., Deng, C., Brown, J. A., De Giacomo, A. F., Gaw, A. F., Shi, G., Le, Q. T., Brown, J. M., and Koong, A. C. (2005) Identification of mitogen-activated protein kinase signaling pathways that confer resistance to endoplasmic reticulum stress in Saccharomyces cerevisiae. Mol Cancer Res 3, 669–677

47. Bonilla, M., and Cunningham, K. W. (2003) Mitogen-activated protein kinase stimulation of Ca(2+) signaling is required for survival of endoplasmic reticulum stress in yeast. Mol Biol Cell 14, 4296–4305

48. Rousseau, A., and Bertolotti, A. (2016) An evolutionarily conserved pathway controls proteasome homeostasis. Nature 536, 184–189

49. Uppala, J. K., Sathe, L., Chakraborty, A., Bhattacharjee, S., Pulvino, A. T., and Dey, M. (2022) The cap-proximal RNA secondary structure inhibits preinitiation complex formation on HAC1 mRNA. J Biol Chem 298, 101648

50. Kinoshita, T., Yoshida, I., Nakae, S., Okita, K., Gouda, M., Matsubara, M., Yokota, K., Ishiguro, H., and Tada, T. (2008) Crystal structure of human mono-phosphorylated ERK1 at Tyr204. Biochem Biophys Res Commun 377, 1123–1127

51. Manning, G., Whyte, D. B., Martinez, R., Hunter, T., and Sudarsanam, S. (2002) The protein kinase complement of the human genome. Science 298, 1912–1934

52. Kasler, H. G., Victoria, J., Duramad, O., and Winoto, A. (2000) ERK5 is a novel type of mitogen-activated protein kinase containing a transcriptional activation domain. Mol Cell Biol 20, 8382–8389

53. Gotoh, Y., Nishida, E., Shimanuki, M., Toda, T., Imai, Y., and Yamamoto, M. (1993) Schizosaccharomyces pombe Spk1 is a tyrosine-phosphorylated protein functionally related to Xenopus mitogen-activated protein kinase. Mol Cell Biol 13, 6427–6434

54. Ketela, T., Green, R., and Bussey, H. (1999) Saccharomyces cerevisiae mid2p is a potential cell wall stress sensor and upstream activator of the PKC1-MPK1 cell integrity pathway. J Bacteriol 181, 3330–3340

55. Martin, H., Arroyo, J., Sanchez, M., Molina, M., and Nombela, C. (1993) Activity of the yeast MAP kinase homologue Slt2 is critically required for cell integrity at 37 degrees C. Mol Gen Genet 241, 177–184

56. Harrison, J. C., Bardes, E. S., Ohya, Y., and Lew, D. J. (2001) A role for the Pkc1p/Mpk1p kinase cascade in the morphogenesis checkpoint. Nat Cell Biol 3, 417–420

57. Philip, B., and Levin, D. E. (2001) Wsc1 and Mid2 are cell surface sensors for cell wall integrity signaling that act through Rom2, a guanine nucleotide exchange factor for Rho1. Mol Cell Biol 21, 271–280

58. Heinisch, J. J., Lorberg, A., Schmitz, H. P., and Jacoby, J. J. (1999) The protein kinase C-mediated MAP kinase pathway involved in the maintenance of cellular integrity in Saccharomyces cerevisiae. Mol Microbiol 32, 671–680

59. Kaganovich, D., Kopito, R., and Frydman, J. (2008) Misfolded proteins partition between two distinct quality control compartments. Nature 454, 1088–1095

60. Pina, F. J., and Niwa, M. (2015) The ER Stress Surveillance (ERSU) pathway regulates daughter cell ER protein aggregate inheritance. Elife 4

61. Dodou, E., and Treisman, R. (1997) The Saccharomyces cerevisiae MADS-box transcription factor Rlm1 is a target for the Mpk1 mitogen-activated protein kinase pathway. Mol Cell Biol 17, 1848–1859

62. Madden, K., Sheu, Y. J., Baetz, K., Andrews, B., and Snyder, M. (1997) SBF cell cycle regulator as a target of the yeast PKC-MAP kinase pathway. Science 275, 1781–1784

63. Mascaraque, V., Hernaez, M. L., Jimenez-Sanchez, M., Hansen, R., Gil, C., Martin, H., Cid, V. J., and Molina, M. (2013) Phosphoproteomic analysis of protein kinase C signaling in Saccharomyces cerevisiae reveals Slt2 mitogen-activated protein kinase (MAPK)-dependent phosphorylation of eisosome core components. Mol Cell Proteomics 12, 557–574

64. Alonso-Rodriguez, E., Fernandez-Pinar, P., Sacristan-Reviriego, A., Molina, M., and Martin, H. (2016) An Analog-sensitive Version of the Protein Kinase Slt2 Allows Identification of Novel Targets of the Yeast Cell Wall Integrity Pathway. J Biol Chem 291, 5461–5472

65. Ray, A., Hector, R. E., Roy, N., Song, J. H., Berkner, K. L., and Runge, K. W. (2003) Sir3p phosphorylation by the Slt2p pathway effects redistribution of silencing function and shortened lifespan. Nat Genet 33, 522–526

66. Soulard, A., Cremonesi, A., Moes, S., Schutz, F., Jeno, P., and Hall, M. N. (2010) The rapamycin-sensitive phosphoproteome reveals that TOR controls protein kinase A toward some but not all substrates. Mol Biol Cell 21, 3475–3486

67. Jin, C., Strich, R., and Cooper, K. F. (2014) Slt2p phosphorylation induces cyclin C nuclear-to-cytoplasmic translocation in response to oxidative stress. Mol Biol Cell 25, 1396–1407

68. Alonso-Rodriguez, E., Fernandez-Pinar, P., Sacristan-Reviriego, A., Molina, M., and Martin, H. (2016) An Analog-sensitive Version of the Protein Kinase Slt2 Allows Identification of Novel Targets of the Yeast Cell Wall Integrity Pathway. J Biol Chem 291, 5461–5472

69. Yurko, N., Liu, X., Yamazaki, T., Hoque, M., Tian, B., and Manley, J. L. (2017) MPK1/SLT2 Links Multiple Stress Responses with Gene Expression in Budding Yeast by Phosphorylating Tyr1 of the RNAP II CTD. Mol Cell 68, 913–925 e913

70. Chao, J. T., Pina, F., and Niwa, M. (2021) Regulation of the early stages of endoplasmic reticulum inheritance during ER stress. Mol Biol Cell 32, 109–119

71. Pellegrini, L., Tan, S., and Richmond, T. J. (1995) Structure of serum response factor core bound to DNA. Nature 376, 490–498

72. Lin, T. A., Kong, X., Haystead, T. A., Pause, A., Belsham, G., Sonenberg, N., and Lawrence, J. C., Jr. (1994) PHAS-I as a link between mitogen-activated protein kinase and translation initiation. Science 266, 653–656

73. Pause, A., Belsham, G. J., Gingras, A. C., Donze, O., Lin, T. A., Lawrence, J. C., Jr., and Sonenberg, N. (1994) Insulin-dependent stimulation of protein synthesis by phosphorylation of a regulator of 5’-cap function. Nature 371, 762–767

74. Castelli, L. M., Talavera, D., Kershaw, C. J., Mohammad-Qureshi, S. S., Costello, J. L., Rowe, W., Sims, P. F., Grant, C. M., Hubbard, S. J., Ashe, M. P., and Pavitt, G. D. (2015) The 4E-BP Caf20p Mediates Both eIF4E-Dependent and Independent Repression of Translation. PLoS Genet 11, e1005233

75. Rutter, J., Probst, B. L., and McKnight, S. L. (2002) Coordinate regulation of sugar flux and translation by PAS kinase. Cell 111, 17–28

76. Huse, M., and Kuriyan, J. (2002) The conformational plasticity of protein kinases. Cell 109, 275–282

77. Yang, L., Zheng, L., Chng, W. J., and Ding, J. L. (2019) Comprehensive Analysis of ERK1/2 Substrates for Potential Combination Immunotherapies. Trends Pharmacol Sci 40, 897–910

78. Busca, R., Pouyssegur, J., and Lenormand, P. (2016) ERK1 and ERK2 Map Kinases: Specific Roles or Functional Redundancy? Front Cell Dev Biol 4, 53

79. Gkouveris, I., and Nikitakis, N. G. (2017) Role of JNK signaling in oral cancer: A mini review. Tumour Biol 39, 1010428317711659

80. Trempolec, N., Dave-Coll, N., and Nebreda, A. R. (2013) SnapShot: p38 MAPK substrates. Cell 152, 924–924 e921

81. Unal, E. B., Uhlitz, F., and Bluthgen, N. (2017) A compendium of ERK targets. FEBS Lett 591, 2607–2615

82. Jacobs, D., Glossip, D., Xing, H., Muslin, A. J., and Kornfeld, K. (1999) Multiple docking sites on substrate proteins form a modular system that mediates recognition by ERK MAP kinase. Genes Dev 13, 163–175

83. Mannan, M. A., Shadrick, W. R., Biener, G., Shin, B. S., Anshu, A., Raicu, V., Frick, D. N., and Dey, M. (2013) An ire1-phk1 chimera reveals a dispensable role of autokinase activity in endoplasmic reticulum stress response. J Mol Biol 425, 2083–2099

84. Schneider, C. A., Rasband, W. S., and Eliceiri, K. W. (2012) NIH Image to ImageJ: 25 years of image analysis. Nat Methods 9, 671–675

